# Functional and antigenic landscape of the Nipah virus receptor binding protein

**DOI:** 10.1101/2024.04.17.589977

**Authors:** Brendan B. Larsen, Teagan McMahon, Jack T. Brown, Zhaoqian Wang, Caelan E. Radford, James E. Crowe, David Veesler, Jesse D. Bloom

**Affiliations:** Basic Sciences Division and Computational Biology Program, Fred Hutch Cancer Center, Seattle, WA 98109, USA; Department of Biochemistry, University of Washington, Seattle, WA 98195, USA; Department of Pathology Microbiology and Immunology, The Vanderbilt Vaccine Center, Department of Pediatrics, Vanderbilt University Medical Center, Nashville, TN 37232, USA; Howard Hughes Medical Institute, Seattle, WA 98195, USA

## Abstract

Nipah virus recurrently spills over to humans, causing fatal infections. The viral receptor-binding protein (RBP or G) attaches to host receptors and is a major target of neutralizing antibodies. Here we use deep mutational scanning to measure how all amino-acid mutations to the RBP affect cell entry, receptor binding, and escape from neutralizing antibodies. We identify functionally constrained regions of the RBP, including sites involved in oligomerization, along with mutations that differentially modulate RBP binding to its two ephrin receptors. We map escape mutations for six anti-RBP antibodies, and find that few antigenic mutations are present in natural Nipah strains. Our findings offer insights into the potential for functional and antigenic evolution of the RBP that can inform the development of antibody therapies and vaccines.

## Introduction

Nipah virus is a zoonotic negative-sense RNA virus that circulates in *Pteropus* bats across SE Asia (*1–4*). Nipah virus and some other viruses in the *Henipavirus* genus spill over into humans and livestock with fatality rates approaching 70% (*5–7*). Nipah virus was first identified during an outbreak in Malaysia in 1998 with subsequent (almost annual) spillovers in Bangladesh and India. These spillovers have sometimes resulted in human-to-human transmission, raising concerns about the potential for a larger outbreak (*8*, *9*). No vaccines or specific therapeutics are approved for use in humans against Nipah virus.

Nipah virus has two different surface proteins that mediate cell entry: the tetrameric receptor-binding protein (RBP or G) and the trimeric fusion (F) protein. RBP binds to the cell-surface proteins ephrin-B2 or -B3 (EFNB2/3), either of which can function as the viral receptor (*10–13*). EFNB2 and EFNB3 are members of a protein family that is crucial for vertebrate development and cell signaling, and are highly conserved among mammals (*14–16*). The Nipah virus RBP binds both EFNB2 and EFNB3 despite these two receptor proteins only sharing 40% amino-acid identity (*12*). The affinity of RBP for EFNB2 is among the highest of any known viral protein for its receptor, while the affinity for EFNB3 is ∼25-fold lower (*13*). Following binding to its receptor(s), the RBP undergoes a conformational shift that triggers F to fuse the viral and cell membranes (*17*).

Potent RBP-directed monoclonal antibodies have been identified that neutralize Nipah virus and prevent disease in animal models (*18–21*). Antibodies and vaccines are currently being developed as a defense against Nipah virus (*22–29*), but for some other viruses, evolution has rendered such countermeasures less effective (*30*). In vitro studies have identified some RBP antibody-escape mutations (*31*, *32*), but such studies have been limited due to the inherent difficulty of working with Nipah virus itself, which is a biosafety level 4 (BSL-4) pathogen. There are also relatively few sequences of natural Nipah viruses, limiting the inferences that can be made about evolutionary constraints from sequence variation.

Here we experimentally measure the effects of all amino-acid mutations to the RBP ectodomain using a BSL-2 lentiviral pseudovirus deep mutational scanning (DMS) platform (*33*, *34*). By coupling experimental selections on variant libraries with deep sequencing, we quantify how mutations affect three RBP phenotypes: cell entry, receptor binding, and antibody escape. Collectively, these results elucidate the evolutionary potential of a key protein from this pathogenic zoonotic virus with pandemic potential.

## Results

### A pseudovirus deep mutational scanning library of the Nipah virus RBP

To measure how mutations to RBP impact cell entry, receptor binding and antibody evasion, we utilized a recently developed DMS platform (*33*, *34*) to create genotype-phenotype-linked libraries of lentiviruses pseudotyped with mutants of RBP alongside the unmutated Nipah F protein (fig S1). The pseudotyped lentiviruses are non-replicative and encode no viral proteins other than the RBP, and so provide a tool to safely study RBP mutants at BSL-2. We mutagenized RBP from the Nipah Malaysia strain, which was the first described strain of the virus and is widely used in other published work. The Nipah Malaysia RBP differs from other known Nipah RBPs by a maximum of 29 amino-acid mutations (fig S2). We truncated 32 and 22 amino acids from the cytoplasmic tails of RBP and F, respectively, which increased pseudovirus titers without apparent effect on RBP antigenicity (*35*, *36*) (fig S3).

We made duplicate mutant libraries targeting all amino-acid mutations of the RBP ectodomain (residues 71 to 602), for a total of 532×19 = 10,108 mutations. Our final duplicate libraries contained 78,450 and 60,623 unique barcoded RBP variants that each covered >99.5% of all possible mutations (fig S4). Most RBP variants carried just a single amino-acid mutation, although there were some variants with multiple mutations (fig S4).

We created target cells that expressed only EFNB2 or EFNB3 to distinguish the effects of RBP mutations on usage of each receptor. To do this, we transduced either EFNB2 or EFNB3 into CHO cells, which do not express any ephrins (*13*) (fig S5). We used EFNB2/3 orthologs from a natural Henipavirus host, the black flying fox (*Pteropus alecto*) (*37*) to avoid the possibility that our experiments could generate potentially hazardous information (*38*, *39*) about mutations that adapted RBP to better bind to human receptors. We refer to these bat orthologs as bEFNB2 or bEFNB3; they share 95% and 97% amino-acid identity with the corresponding human proteins (fig S6). Both bEFNB2- and bEFNB3-expressing CHO cells could be efficiently infected with Nipah RBP pseudovirus (fig S5).

### Effects of RBP mutations on cell entry

We measured the effects of all RBP amino-acid mutations on pseudovirus entry in CHO-bEFNB2 and CHO-bEFNB3 cells by quantifying the ability of each barcoded pseudovirus variant to enter cells when pseudotyped with its RBP mutant and unmutated F versus VSV-G (fig S7A). Because our libraries contain some RBPs encoding multiple mutations (fig S4), we deconvolved the effects of individual mutations using a global epistasis model (*40*). After excluding low-confidence measurements, our dataset contained cell-entry measurements for 97% and 96% of the 10,108 possible single amino-acid mutants of the RBP ectodomain in CHO-bEFNB2 and CHO-bEFNB3 cells, respectively. Measurements of mutational effects on cell entry made using the two independent libraries were highly correlated (fig S8), and throughout this paper we report the average measurement across the two libraries.

Mutations had varied effects on RBP-mediated entry in CHO-bEFNB3 cells, ranging from highly deleterious to well tolerated (Fig 1; see also CHO-bEFNB2 data in fig S9). A cryo-electron microscopy structure of the Nipah virus RBP ectodomain tetramer revealed that it consists of four major regions designated stalk, neck, linker, and head (*31*) (fig S10). Functional constraints were particularly high in the stalk and neck regions, the area of the head interacting with the EFN receptors, and the dimerization interface between distal heads (Fig 2A-D).

**Figure 1.**
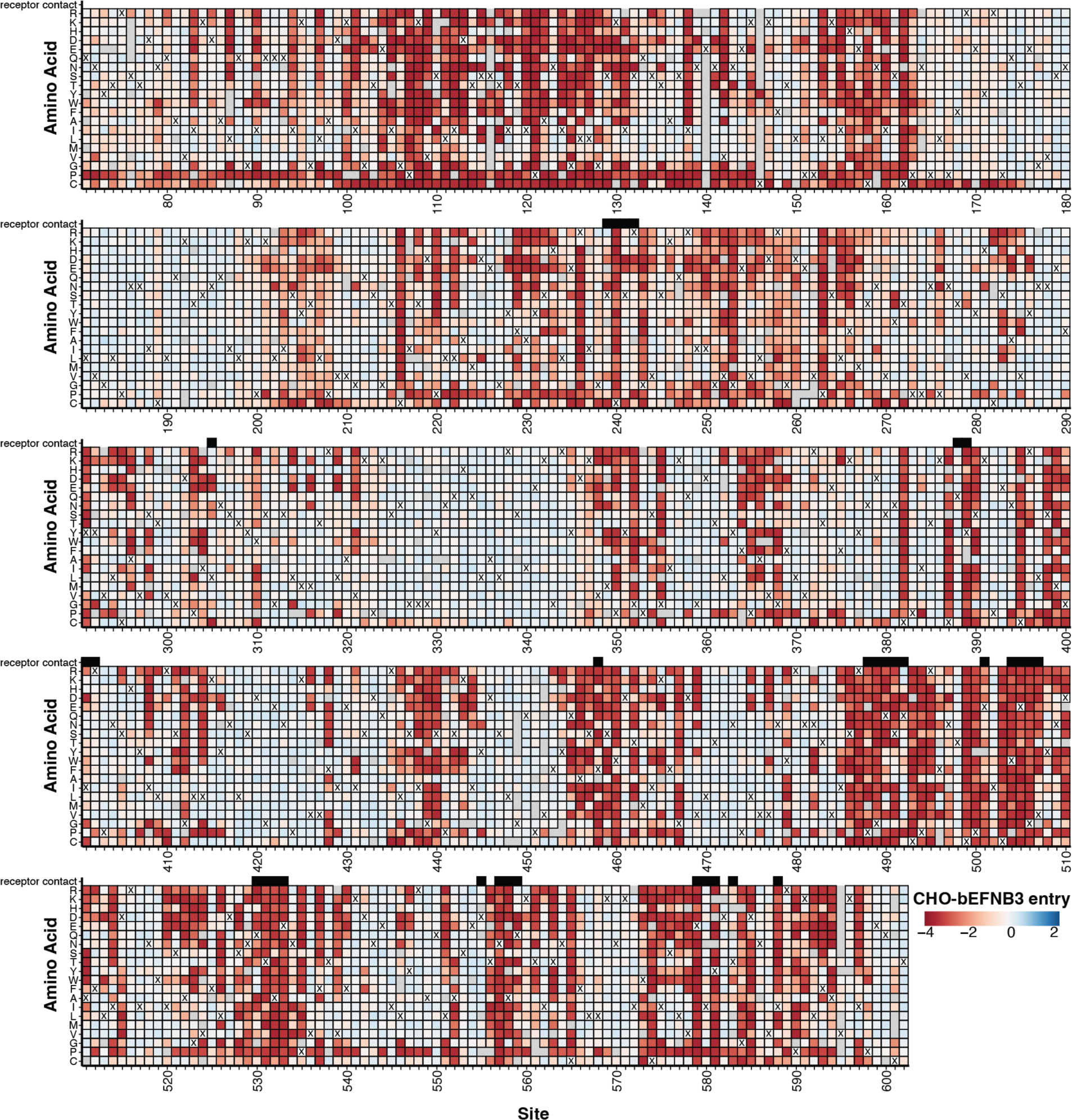
Effects of RBP mutations on entry into CHO-bEFNB3 cells. For each mutation, the entry score reflects the cell entry efficiency of a pseudovirus with that RBP mutation relative to the unmutated RBP. Negative values (red) indicate impaired entry, zero (white) indicates no effect, and positive values (blue) indicate improved entry. The wildtype amino-acid in the Malaysia strain RBP at each site is indicated with a ‘X’. Mutations that were not measured with high confidence in our experiments are indicated with a light gray box. Residues that directly contact the EFNB3 receptor (<= 4 angstroms of the receptor calculated from PDB 3D12) are indicated with black boxes above the heatmap. An interactive version of this heatmap is available at https://dms-vep.org/Nipah_Malaysia_RBP_DMS/htmls/E3_entry_heatmap.html. See fig S9 for comparable data for cell entry into CHO-bEFNB2 cells.

**Figure 2.**
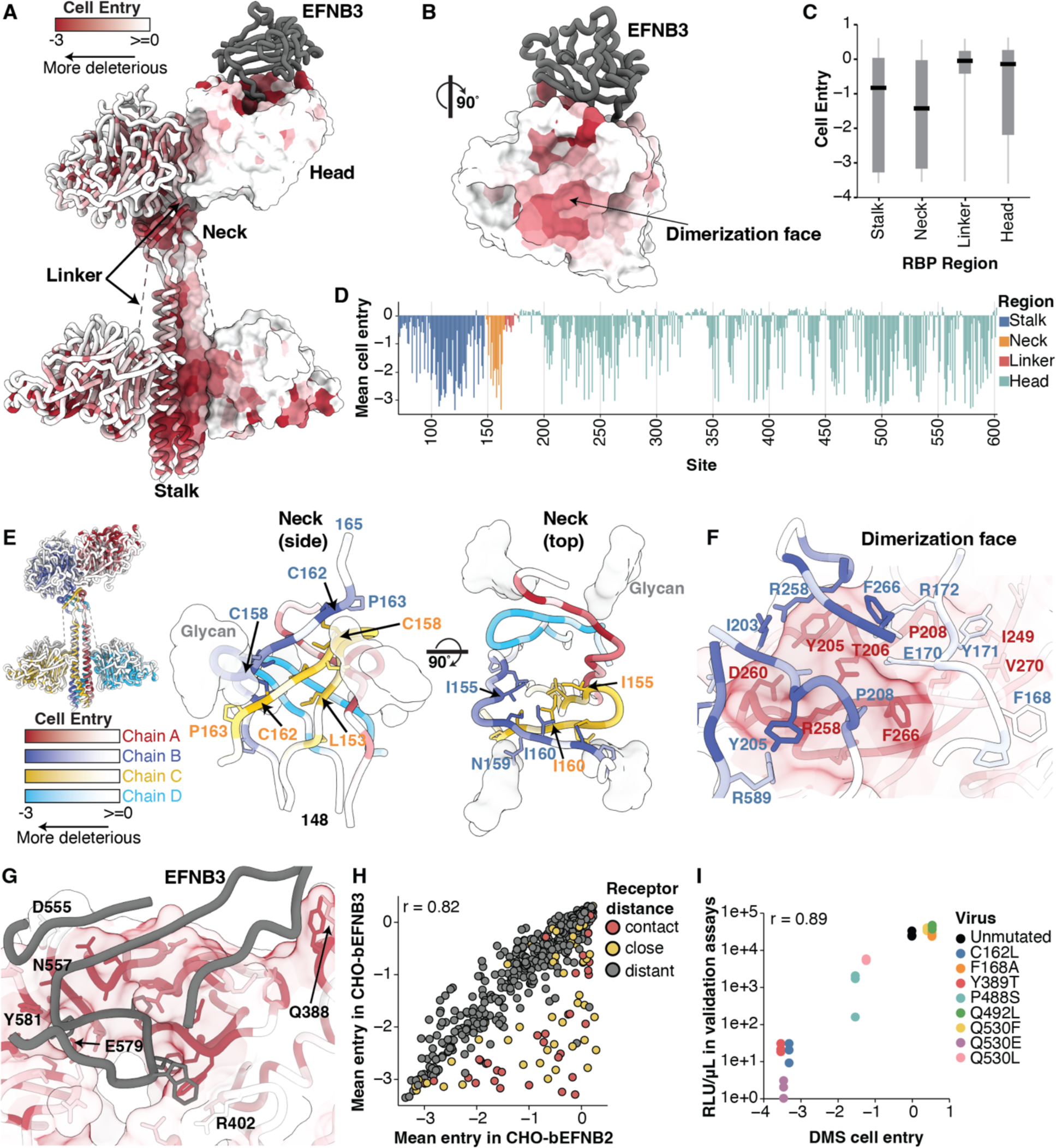
Functional constraint on different regions of RBP for entry into CHO-bEFNB3 cells. A) RBP tetrameric structure colored by the effect of mutations on cell entry (PDBs: 7TXZ, 7TY0, 3D12). Each site is colored by the mean effect of all amino-acid mutations at that site, with red indicating impaired cell entry, and white indicating entry comparable to unmutated RBP. B) Effects of mutations on cell entry projected on the RBP head oriented to visualize the RBP dimerization face. C) Boxplot showing the impacts of mutations on cell entry across different RBP regions. D) Average effects of mutations at each RBP site on cell entry. E) Average effect of mutations at each site on cell entry for the RBP neck viewed from the side or top. Each chain has a unique color scale as indicated in the color-scale bars, with darker colors indicating impaired entry. F) Effects of mutations on entry for the interface between the distal heads (chains A and B). G) Effects of mutations on entry at the RBP/EFNB3 interface. The same color scale is used as in (A). EFNB3 is shown as a gray cartoon. H) Correlation between effects of mutations on cell entry in CHO-bEFNB2 vs CHO-bEFNB3 cells. Each point is the average effect of mutations at a site. Points are colored by distance to the closest receptor residue (contact sites defined as < 4 Å to receptor, close > 4 and < 8 Å, and distant >10 Å). I) Validation assays showing the correlation between single mutant pseudovirus titers in CHO-bEFNB3 cells versus effects measured in DMS. Three independent measurements were made for each validation pseudovirus. Infectivity was quantified by luciferase signal (RLU/µL) at 48 hours after infection of CHO-bEFNB3 cells.

The RBP neck was the most functionally constrained region, probably due to its critical role in tetramerization, stability, and F-triggering (*41*) (Fig 2C). The neck forms a stacked β-sandwich, encircled by glycans linked to N159 (Fig 2E). The cysteines at sites 158 and 162 were intolerant to any mutations, consistent with previous mutagenesis work (*41*) and their key structural role (*31*). The neck is particularly rich in prolines and hydrophobic residues, which were generally functionally constrained (Fig 2E, fig S11A). Previous work suggested the N-linked glycan at position 159 is important for F-triggering (*42*). Our results indicate strong constraint on site 159, as most mutations at this site were highly deleterious. However, mutations to the negatively charged D and E amino acids at site 159 had neutral effects on entry (Fig 1), suggesting the position’s negative charge might be more important for cell entry than the glycan itself.

The distal heads form a complex, asymmetric dimerization interface with multiple interacting residues. Nearly all mutations at sites Y205, T206, P208, R258, G259, and F266 in the dimerization interface drastically reduced cell entry (Fig 2F, Fig 1). At site Y205, the only non-deleterious change was mutation to a tryptophan, another aromatic side chain (Fig 1), likely due to retention of cation-pi and van der Waals interactions between the residue 205 aromatic side chain and R258 from the adjacent head (Fig 2F). Although several sites in the linker (164–173) from chain B contact chain A residues in the dimer interface (Fig 2F), most mutations at linker sites had minor effects on entry, indicating these individual interactions are non-essential (Fig 2F, Fig 1).

Most mutations at sites that directly contact the receptor were highly deleterious for entry in CHO-bEFNB3 cells (Fig 2G). Sites on the periphery of the binding pocket were more tolerant to mutations than those within the central binding pocket (Fig 2G). D555, Q388, and R402 each tolerated several mutations, whereas sites N557, E579 and Y581 were the most constrained of all receptor-interface sites and all mutations were highly deleterious (Fig 2G, fig S11B, fig S12).

The effects of mutations on entry in CHO-bEFNB2 and CHO-bEFNB3 cells were largely similar (Fig 2H). However, some receptor-contact sites were more tolerant to mutations for entry into CHO-bEFNB2 than CHO-bEFNB3 cells (Fig 2H, fig S12). This difference is likely because RBP has a ∼25-fold higher binding affinity for EFNB2 compared to EFNB3, meaning it can tolerate more reduction in binding while still maintaining efficient entry in bEFNB2-expressing cells, at least in the cell-culture context of our experiments (*13*).

To validate the DMS measurements of mutational effects on cell entry, we constructed eight pseudoviruses with single RBP mutations spanning a range of entry effects from DMS, and evaluated their entry in CHO-bEFNB2 and CHO-bEFNB3 cells relative to unmutated RBP. Cell entry for these individual pseudoviruses strongly correlated with the DMS measurements (r=0.89 for CHO-bEFNB3, Fig 2I; r=0.79 for CHO-bEFNB2, fig S13).

### Effects of RBP mutations on receptor binding

Mutations to RBP can affect cell entry by many mechanisms, including altering receptor binding, fusion triggering, protein stability, or protein expression. To partially deconvolve these mechanisms, we sought to measure the effects of RBP mutations on binding to bEFNB2 and bEFNB3. Inspired by prior studies that show inhibition of viral entry by soluble receptor is proportional to receptor binding affinity (*43*, *44*), we incubated our RBP pseudovirus libraries with different concentrations of soluble bEFNB2 or bEFNB3 to measure how RBP mutations affect receptor binding (fig S7B). This approach measures the effects on binding for mutations that support at least moderate cell entry; highly deleterious mutations that completely disrupt entry will not yield infectious pseudovirus.

We first tested neutralization of pseudovirus encoding unmutated RBP by soluble bEFNB2 or bEFNB3. Soluble monomeric and dimeric bENFB2 potently neutralized pseudovirus with half-maximal inhibitory concentrations (IC_50_) of 0.9 or 0.05 nM, respectively. In contrast, monomeric or dimeric soluble bEFNB3 had IC_50_s of >400 or 0.6 nM (Fig 3A). The higher potency of bEFNB2 was expected, given the higher binding affinity of the RBP for EFNB2 relative to EFNB3. We chose to use monomeric bEFNB2 and dimeric bEFNB3 for our DMS library selections due to their similar neutralizing potencies. In total, we measured the effects on binding for 6,672 RBP mutations for bEFNB2 and 6,503 mutations for bEFNB3 (fig S14,15). Most mutations for which binding was not measured were highly deleterious for pseudovirus entry (fig S16).

**Figure 3.**
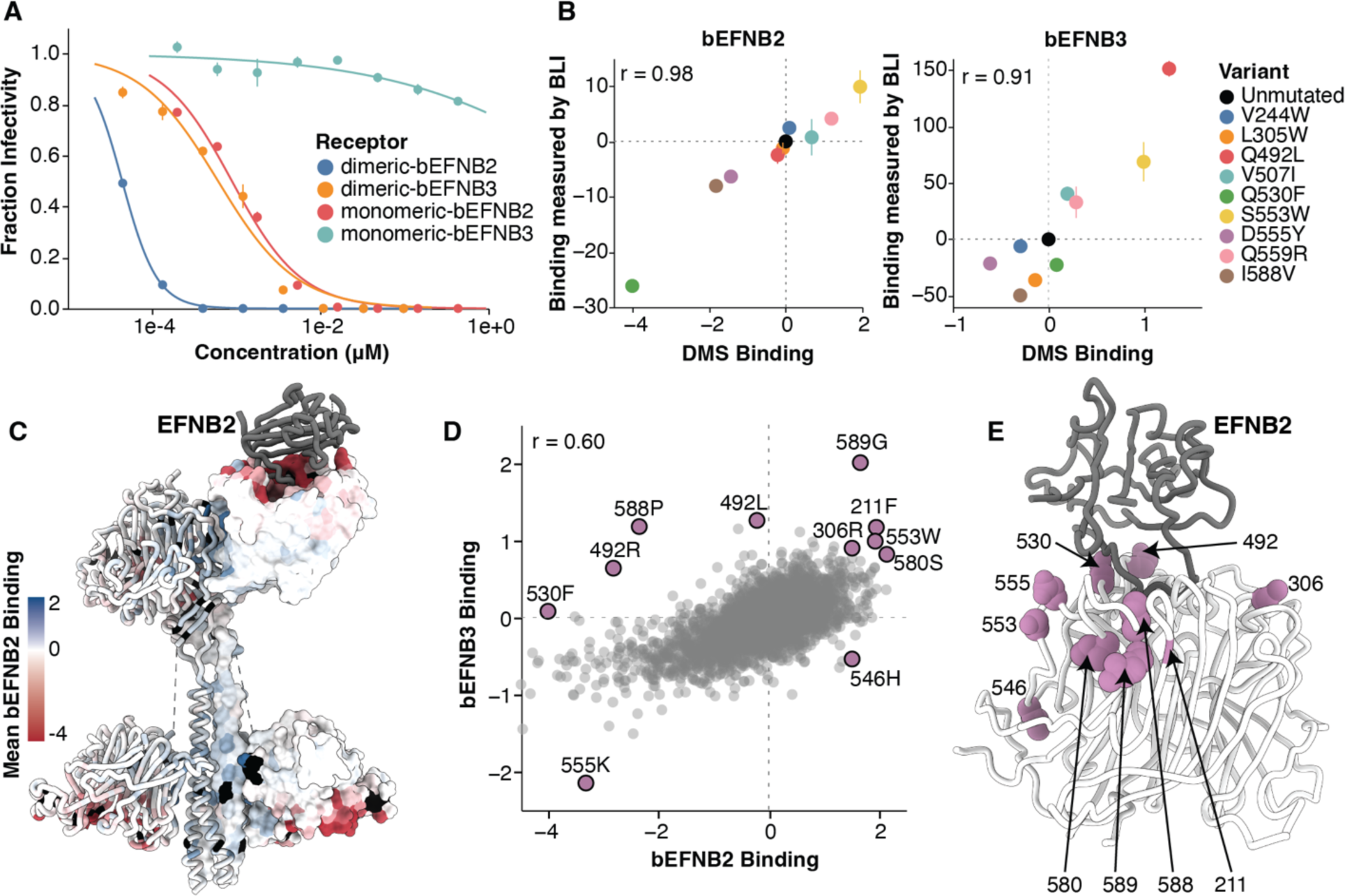
Effects of RBP mutations on binding to bEFNB2 or bEFNB3. A) Neutralization of unmutated Nipah RBP/F pseudoviruses by monomeric or dimeric soluble bEFNB2 or bEFNB3. B) Correlation between biolayer interferometry (BLI) measurements of binding affinity with DMS measured effects of mutations on binding. The magnitude of binding was assessed by quantifying the change in total area under the curve (AUC) relative to unmutated RBP for binding of the indicated RBP head domains to immobilized dimeric bEFNB2 or bEFNB3 (see fig S18 for raw BLI sensorgrams). C) Tetrameric structure of RBP colored by the site-average effects of mutations on binding to bEFNB2. Sites that are missing binding measurements (typically because mutations are highly deleterious for cell entry) are colored black. D) Correlation between the effects of mutations on binding to bEFNB2 versus bEFNB3. Mutations with notable effects on binding are labeled. E) Location of sites on RBP’s head that have notable mutations on binding. For heatmaps showing effects of all mutations on binding as measured by DMS, see fig S14,15.

To validate that these DMS measurements indeed reflected the strength of RBP binding to bEFNB2/3, we produced the soluble monomeric head domains for nine RBP variants that spanned a range of binding effects from DMS (fig S17), and quantified their binding to immobilized bEFNB2 and bEFNB3 using biolayer interferometry (BLI) (Fig 3B, fig S18). The BLI binding measurements were highly correlated with the DMS measurements (r=0.98 for bEFNB2; r=0.91 for bEFNB3; Fig 3B). The RBP mutations tested by BLI did not affect binding to a monoclonal antibody (HENV-32, which recognizes a conformational epitope (*20*)), demonstrating they are specifically altering receptor binding and not proper head folding (fig S19). We also validated that neutralization of six single RBP mutant pseudoviruses by soluble bEFNB2 and bEFNB3 correlated well with the DMS (fig S20).

Most mutations at receptor contact sites greatly reduced binding to both bEFNB2 and bEFNB3, although mutations across the rest of the RBP had largely neutral effects on binding (Fig 3C,D; fig S14,15,21). Mutations that had opposite effects on binding to bEFNB2 and bEFNB3 were primarily located near the receptor interface (Fig 3D, fig S22). Several mutations at site 492 increased binding to bEFNB3 while decreasing binding to bEFNB2 (Fig 3D, Fig 3B, fig S17, fig S23). Site 492 is in close proximity to three amino acids in the receptor that differ between bEFNB2 and bEFNB3, and one site that differs between human and bat EFNB2 (fig S23). Several mutations in the loop containing sites 580-590 also had large effects on binding to bEFNB2 versus bEFNB3 (Fig 3D, fig S24)

Mutations that caused the largest increases in receptor binding were scattered at sites throughout the head (Fig. 3E). For instance, S553W increases binding to both bEFNB2 and bEFNB3 even though it does not contact any receptor residues (Fig 3E, Fig 3B, fig S22). This mutation may act by shifting the position of the 580-590 loop, or the receptor contact sites 557-559. Several mutations at site N306, which is glycosylated, also increased binding to both bEFNB2 and bEFNB3 (fig S22). These mutations remove the glycan, likely reducing possible steric hindrance of some of the oligosaccharide orientations with receptor engagement.

### Effects of RBP mutations on antibody escape

We sought to prospectively map how all mutations affected neutralization by six antibodies targeting RBP to define potential escape sites, and understand if escape mutations are already circulating or are likely to be tolerated in future evolution. To make these measurements, we incubated the pseudovirus libraries with different concentrations of each antibody, followed by infection of CHO-bEFNB3 target cells, and then deep sequencing to quantify the infection by each RBP mutant relative to a non-neutralizable standard unaffected by the antibody (fig S7C).

We chose potent neutralizing antibodies that target a diversity of epitopes on the RBP head based on prior structural work or competition mapping studies (*20*, *21*, *31*, *45*) (fig S25). We measured the IC_50_ for each antibody against unmutated RBP/F pseudovirus, which ranged from 12 to 143 ng/mL (fig S25) consistent with previous studies using authentic virus or a replicating Cedar virus chimera (*20*, *21*, *31*, *32*).

Some antibodies were escaped by numerous well-tolerated RBP mutations, whereas others were mostly escaped by mutations that substantially impaired RBP-mediated cell entry (Fig 4A). For example, HENV-103, a dimerization-face-targeting antibody, was escaped primarily at five sites, and most escape mutations at these sites substantially impaired cell entry (Fig 4A). In contrast, nAH1.3 was escaped by mutations at many sites that were well tolerated for cell entry (Fig 4A). Similar trends hold if we examine only amino-acid mutations accessible through a single-nucleotide change to the Malaysia strain Nipah RBP gene sequence (fig S26). These results have implications for antibody countermeasures: for instance, our data suggest there could be a low barrier to resistance to nAH1.3, whereas resistance to HENV-103 may impose a substantial cost on RBP’s essential cell-entry function.

**Figure 4.**
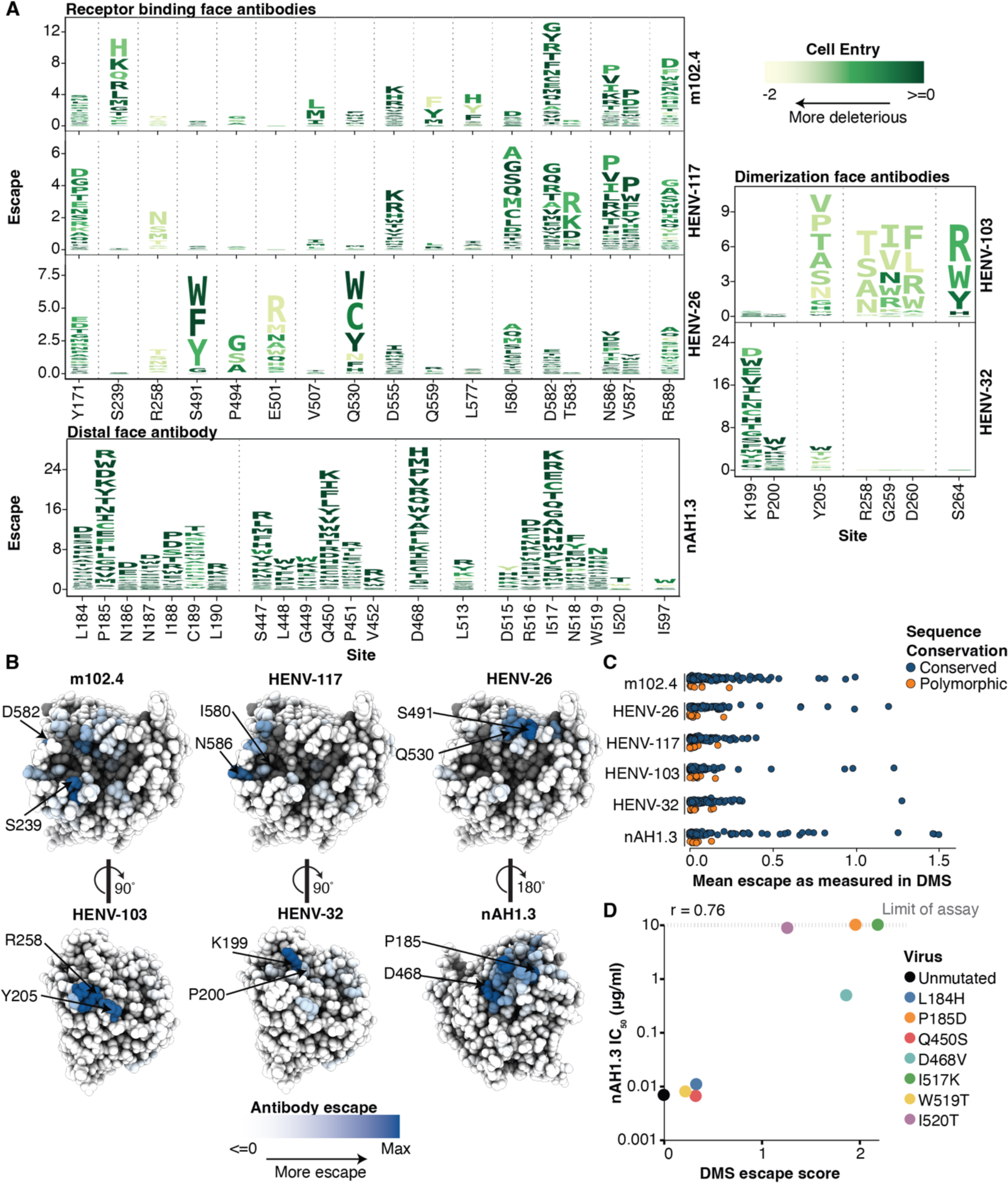
DMS maps of escape mutations from six monoclonal antibodies. A) Key sites of escape for each antibody. The height of each letter is proportional to the escape caused by that amino-acid mutation, and letters are colored by the effect of that mutation on entry in CHO-bEFNB3 cells (dark green indicates well tolerated mutations, light yellow indicates impaired cell entry). These logo plots show the top escape sites for each antibody; for all sites see (https://dms-vep.org/Nipah_Malaysia_RBP_DMS/htmls/mab_plot_all.html). For panels A-C, only mutations that decrease in antibody neutralization are shown. B) Antibody escape mapped onto the structure of RBP’s head. The escape averaged across all mutations at each site is shown as a white-blue color scale, with sites where mutations cause strong escape in darker blue, and those not affecting neutralization in white. Dark gray sites have no escape measurements due to mutations strongly impairing cell entry. C) Variability of RBP among known Nipah virus sequences at sites of antibody escape mapped in this study. Sites are classified as conserved or polymorphic based on whether they show variation among at least two Nipah RBP sequences. D) Correlation between the IC_50_ measured for single mutant Nipah RBP pseudoviruses for antibody nAH1.3 in validation assays versus escape measured in DMS. The upper limit of IC_50_ values in the validation assays was 10 μg/mL (dashed line). See fig S26 for escape logo plots that show only mutations that are single-nucleotide accessible.

The sites of escape for each antibody clustered on the surface of RBP (Fig 4B), near the antibody-binding footprints (fig S27). Antibodies m102.4, HENV-26, and HENV-117 all target the receptor-binding face, but were mostly escaped by distinct mutations (Fig 4A,B). These mutations are at sites mainly along the rim of the receptor pocket, as mutations within the central pocket are likely too deleterious for cell entry to yield functional pseudovirus (Fig 4B; see mutational constraint within the receptor binding pocket in Fig 2G). The two dimerization-face targeting antibodies also had slightly different sites of major escape: HENV-103 was escaped primarily at site 205, while HENV-32 was escaped by almost every mutation at site 199, with only minor escape at sites 200 and 205 (Fig 4A, B).

Mutations at a subset of sites had opposite effects on antibody neutralization for receptor-binding and dimerization-face targeting antibodies (fig S28). For example, mutations at sites 165-172 in the linker region escaped neutralization by receptor binding face targeting antibodies, while increasing neutralization by dimerization face targeting antibodies. These mutations likely alter the conformation of the RBP, which modifies the relative accessibility of each epitope.

Few of the escape mutations identified in our DMS are found in natural RBP sequences (Fig 4C), consistent with the low diversity of known Nipah RBP sequences. The only notable sequence variation was at sites 172 (escapes receptor-binding face antibodies) and 274 (escapes dimerization-face antibodies), which differ between the two major clades of Nipah virus (Malaysian and Bangladesh; fig S2) and could partially explain previously observed differences in certain antibody neutralization potencies between the two strains (*20*, *21*).

To validate the DMS antibody-escape maps, we tested neutralization of seven RBP single mutants in traditional pseudovirus neutralization assays against nAH1.3. The changes in IC_50_ measured in the neutralization assays were highly correlated with the DMS measurements (r=0.76; Fig 4D, fig S29). Escape mutations previously identified using authentic Nipah virus or replicating Cedar virus chimeras also correspond well with our DMS (*31*) (e.g. I520T introduces an N-linked glycosylation site at position N518 of NiV RBP within the nAH.1.3 epitope, fig S30).

## Discussion

We have measured how all mutations to the Nipah virus RBP affect cell entry, receptor binding, and antibody escape. The resulting maps of mutational effects have important implications both for understanding of Nipah virus cell entry and the design of countermeasures. One major unanswered question is how RBP triggers F following receptor binding. RBP undergoes a suite of conformational changes, including exposure of the dimeric interface and propagation of signal from RBP’s head to the neck and stalk, which triggers F (*46*, *47*). Our DMS revealed sites in the dimer interface where mutations strongly impair cell entry (e.g., R258 and F266) that may be involved in the conformational changes that lead to triggering of F by RBP. These results highlight the value of measuring the effects of all mutations in an experimental system that recapitulates the full RBP and F mediated entry process.

Receptor binding affects both the host range and infectivity of viruses (*48*, *49*). RBP is unusual in strongly binding to two different receptors conserved across mammals. EFNB2 is broadly expressed in many tissues, while EFNB3 is expressed largely in the brain (*12*, *50*). EFNB2 is likely the key receptor for Nipah virus for transmission, while EFNB3-mediated infection may contribute to the fatal encephalitis and brain pathology frequently observed in human infections (*12*). Our DMS identified mutations that alter binding to one versus the other bat versions of these receptors, as well as mutations that increase binding to both bat receptors.

Antibody therapies against emerging RNA viruses are a powerful approach for treating spillover infections. However, single mutations are often sufficient to escape neutralization, and can emerge rapidly at functionally unconstrained sites (*51*, *52*). Our work prospectively maps all mutations that escape antibodies, while also assessing the impact of those mutations on RBP function, thereby providing a way to assess the potential for resistance mutations to emerge. There was wide variation in how well the mutations that escape different antibodies were tolerated for RBP function. Some antibodies that targeted either the RBP’s dimerization or receptor-binding face tended to be escaped mostly by mutations that substantially impaired RBP-mediated cell entry, whereas an antibody targeting the RBP’s distal face was escaped by numerous well-tolerated mutations.

Overall, our work defines the functional and antigenic landscape of the Nipah RBP. By measuring how all RBP mutations affect three key protein phenotypes, we define both constraint and plasticity in a way that both informs basic understanding and countermeasures.

## Acknowledgements

We thank Bernadeta Dadonaite for helpful advice during the course of this project. We also thank Stacey Rutherford at Vanderbilt for technical support in monoclonal antibody production. This research was also supported by the Genomics & Bioinformatics Shared Resource, <u>RRID:SCR_022606</u>, of the Fred Hutch/University of Washington/Seattle Children’s Cancer Consortium (P30 CA015704) and the Flow Cytometry Shared Resource, <u>RRID:SCR_022613</u>, of the Fred Hutch/University of Washington/Seattle Children’s Cancer Consortium (P30 CA015704), and by Fred Hutch Scientific Computing, NIH grants S10-OD-020069 and S10-OD-028685.

## Funding

This work was supported in part by the NIH/NIAID under grant R01AI141707 to JDB. as well as DP1AI158186, 75N93022C00036 and U19AI142764 to DV, and a project in grant U19 AI142764 to J.E.C and an Investigators in the Pathogenesis of Infectious Disease Awards from the Burroughs Wellcome Fund (DV), the University of Washington Arnold and Mabel Beckman cryoEM center and the National Institute of Health grant S10OD032290 (to DV). JDB and DV are investigators of the Howard Hughes Medical Institute and DV is the Hans Neurath Endowed Chair in Biochemistry at the University of Washington. BBL is a Washington Research Foundation Postdoctoral Fellow.

## Competing Interests

JDB is on the scientific advisory boards of Apriori Bio, Invivyd, Aerium Therapeutics, and the Vaccine Company. JDB consults for GlaxoSmithKline. JDB receives royalty payments as an inventor on Fred Hutch licensed patents related to viral deep mutational scanning. JEC. has served as a consultant for Luna Labs USA, Merck Sharp & Dohme Corporation, Emergent Biosolutions, GlaxoSmithKline and BTG International Inc. JE. is a member of the scientific advisory board of Meissa Vaccines, a former member of the Scientific Advisory Board of Gigagen (Grifols) and is founder of IDBiologics. The laboratory of JEC received unrelated sponsored research agreements from AstraZeneca, Takeda, and IDBiologics during the conduct of the study. Vanderbilt University has applied for a patent pertinent to some of the materials in this paper.

## Author Contributions

Conceptualization: BBL, JDB

Data curation: BBL, DV, JDB

Formal analysis: BBL, DV, JDB

Funding acquisition: JDB, DV

Investigation: BBL, TM, ZW, JTB

Methodology: JDB, BBL, TM, CER

Project administration: JDB, DV

Resources: DV, ZW, JTB, JEC Jr

Software: JDB, BBL

Supervision: JDB, DV

Validation: BBL, TM

Visualization: BBL, JDB

Writing - original draft: BBL, JDB

Writing - review and editing: JDB, BBL, DV

## Data and Materials Availability

All raw data and code used for data analysis are publicly available here (https://github.com/dms-vep/Nipah_Malaysia_RBP_DMS). A landing page with links to jupyter notebooks, raw data, and interactive visualizations is available here https://dms-vep.org/Nipah_Malaysia_RBP_DMS/). Additional information is available in supplementary materials.

## Methods

### Data availability and interactive plots of results

All code and data are publicly available here (https://github.com/dms-vep/Nipah_Malaysia_RBP_DMS), with a landing page with links to specific Jupyter notebooks used at each step in the analysis (https://dms-vep.org/Nipah_Malaysia_RBP_DMS/). The main analyses were all performed with dms-vep-pipeline-3 (https://github.com/dms-vep/dms-vep-pipeline-3).

Raw data from all DMS experiments are available in the main GitHub repository (https://github.com/dms-vep/Nipah_Malaysia_RBP_DMS), with final filtered .csv files in this directory (https://github.com/dms-vep/Nipah_Malaysia_RBP_DMS/tree/master/results/filtered_data/public_filtered)

To explore the cell entry data, heatmaps are available separately for CHO-bEFNB2 (https://dms-vep.org/Nipah_Malaysia_RBP_DMS/htmls/E2_entry_heatmap.html) and CHO-bEFNB3 (https://dms-vep.org/Nipah_Malaysia_RBP_DMS/htmls/E3_entry_heatmap.html) which provide the data with our recommended filters.

If further exploration of the data is desired, there are also heatmaps which provide greater control of parameter filtering (CHO-bEFNB2 here: https://dms-vep.org/Nipah_Malaysia_RBP_DMS/htmls/CHO_bEFNB2_func_effects.html, CHO-bEFNB3 here: https://dms-vep.org/Nipah_Malaysia_RBP_DMS/htmls/CHO_bEFNB3_func_effects.html).

The key filtering parameters for these plots are described below:

- *minimum times_seen*: The minimum number of times a mutation is detected, averaged across libraries and replicates. Mutations with higher ‘times_seen’ are more likely to have more accurate measurements of effects.
- *maximum effect_std*: The maximum cutoff for the standard deviation of mutant effects. Mutations with lower standard deviations are more likely to be accurate.
- *minimum n_selections*: The minimum number of independent selections a mutation must be detected.
- *minimum max of effect at site*: Filters sites based on the minimum max effect. Allows the users to only view a subset of sites which have beneficial mutations.

To explore the receptor binding data, interactive heatmaps are available for bEFNB2 (https://dms-vep.org/Nipah_Malaysia_RBP_DMS/htmls/E2_binding_heatmap.html) and bEFNB3 (https://dms-vep.org/Nipah_Malaysia_RBP_DMS/htmls/E3_binding_heatmap.html). These have been pre-filtered based on specific criteria (see methods section *Data filtering*) and represent the easiest way to explore the data. For more control over the heatmaps, including parameter filtering and additional interactivity, pages are available here for bEFNB2 (https://dms-vep.org/Nipah_Malaysia_RBP_DMS/htmls/bEFNB2_monomeric_mut_effect.html) and bEFNB3 (https://dms-vep.org/Nipah_Malaysia_RBP_DMS/htmls/bEFNB3_dimeric_mut_effect.html). Here, users can also use the sliders described previously, along with adjusting the *minimum functional effect*, which controls the minimum cell entry score for a mutant to be shown in the receptor binding heatmap.

To explore the antibody escape data, an interactive heatmap for all antibodies is here (https://dms-vep.org/Nipah_Malaysia_RBP_DMS/htmls/mab_plot_all.html). The data have been pre-filtered and enable easier interpretation, and includes information about antibody contact distance for the antibodies that have structures available. More in-depth heatmaps are available for each antibody separately:

- m102.4 (https://dms-vep.org/Nipah_Malaysia_RBP_DMS/htmls/m102.4_mut_effect.html)
- nAH1.3 (https://dms-vep.org/Nipah_Malaysia_RBP_DMS/htmls/nAH1.3_mut_effect.html)
- HENV-26 (https://dms-vep.org/Nipah_Malaysia_RBP_DMS/htmls/HENV26_mut_effect.html)
- HENV-32 (https://dms-vep.org/Nipah_Malaysia_RBP_DMS/htmls/HENV32_mut_effect.html)
- HENV-103 (https://dms-vep.org/Nipah_Malaysia_RBP_DMS/htmls/HENV103_mut_effect.html)
- HENV-117 (https://dms-vep.org/Nipah_Malaysia_RBP_DMS/htmls/HENV117_mut_effect.html)

These heatmaps give the user further control over the filtering parameters with interactive sliders.

### Plasmid maps and primer sequences

Maps for all plasmids used in study are available here (https://github.com/dms-vep/Nipah_Malaysia_RBP_DMS/tree/master/data/custom_analyses_data/paper_supplementary_data/plasmid_maps). Primer sequences are available here (https://github.com/dms-vep/Nipah_Malaysia_RBP_DMS/blob/master/data/custom_analyses_data/paper_supplementary_data/primer_sequences.csv).

### Creation of CHO-bEFNB2 and CHO-bEFNB3 target cells

In order to isolate the effects of EFNB2 and EFNB3 on viral entry, we sought to make stable cells that would only express one receptor at a time. In addition, to alleviate biosecurity concerns with identifying human specific adaptations, we made target cells that express the bat orthologs of each receptor, rather than human. Finally, we wanted the expression of bEFNB2 and bEFNB3 in the target cells to be roughly similar to a clonal cell line (293T), as extremely high expression of ephrin could bias the DMS results (*53*).

To that end, we used Chinese Hamster Ovary (CHO pgsA-745) cells from ATCC (CRL-2242) because they do not express any ephrins (*11*, *12*). Next, we identified the EFNB2 (GenBank Accession: NP_001277099.1) and EFNB3 (ELK03828.1) amino-acid sequences from a natural host of Henipaviruses (*Pteropus alecto*, the black flying fox) in GenBank. Codon optimized versions of these sequences were synthesized by Twist Biosciences. These sequences were cloned into a lentivirus backbone (see the maps for the plasmids 3949_pHAGE2_EF1A_BatEphrinB2_T2A_mCherry_CMV_PuroR_vlow, 3694_pHAGE2_EF1A_IRES_mCherry_Pteropus_alecto_EFNB3_HA_tag_Extracellular linked in the *Plasmid maps* section of the methods above) under the EF1 alpha promoter. The bEFNB3 construct included an N-terminal extracellular HA tag while the bEFNB2 construct did not. The reason the motifs flanking the bEFNB2 and bEFNB3 sequences are slightly different is because we originally used FLAG and HA tags on both bEFNB2 and bEFNB3, respectively, and attempted to sort cells with different expression levels based on staining with anti-FLAG and anti-HA antibodies. While the bEFNB3 clones had reasonable expression levels as measured by flow cytometry, all of the bEFNB2 clones had very high levels of expression. Therefore, we modified bEFNB2 expression further by introducing a non-ideal 6bp Kozak sequence (AATTTT) before the bEFNB2 start codon (*53*). This resulted in isolation of a CHO-bEFNB2 clone that had lower bEFNB2 expression than the previous attempt, which was comparable to ephrin expression in 293T cells and the bEFNB3 clones (<u>fig S5</u>). Specific details about stable cell isolation are given below.

To produce lentivirus expressing bEFNB2 or bEFNB3, we transfected 1 ug of backbone with 250 ng of lentiviral helper plasmids (26_HDM_Hgpm227, 27_HDM_tat1b, 28_pRC_CMV_Rev1b, 29_HDM_VSV_G) into 293T cells using the BioT transfection reagent following manufacturers directions. After 48 hours, the supernatant was passed through a 0.45 µm syringe filter (Corning, Cat. No. 431220) and stored at −80°C. The generated lentivirus was then used to infect CHO cells at an MOI between 0.1 and 0.01. Individual bEFNB3 expressing cells were sorted 48 hrs later into individual wells of a 96 well plate on a BD Aria II and gated to include only live, single, mCherry positive cells. bEFNB2 expressing CHO cells were passaged in puromycin (1-5 ug/mL) for 1 week and then stained with EphB3-Fc (5667-B3-050; R&D Systems), a natural ligand of EFNB2/3, followed by secondary staining with a Goat anti-human IgG Fc (ThermoFisher A18818). Individual bEFNB2 expressing cells were then sorted into individual wells of a 96 well plate on a Sony MA900 and gated to include only live, single, mCherry positive cells with EFNB expression (FITC) in the same range of 293T cells. Clones of both cell types were then selected based on additional EphB3 staining to ensure they were expressing correct levels of either bEFNB2 or bEFNB3 and for their ability to support high Nipah pseudotyped lentivirus titers (<u>fig S5</u>).

### Anti-RBP monoclonal antibodies

m102.4 was synthesized by GenScript based on the original sequence deposited of the antibody (*45*, *54*). HENV-26, HENV-32, HENV-103, HENV-117, and nAH1.3 were produced as previously described (*20*, *21*, *31*)

### Lentivirus backbone for RBP deep mutational scanning

The lentivirus vector we used for the pseudovirus deep mutational scanning has been previously described (*33*, *34*), and is schematized in <u>fig S1A</u>. A codon-optimized Nipah RBP sequence based on GenBank accession NC_002728.1 lacking 33 amino acids in the cytoplasmic tail at the N-terminus of the protein was inserted into a pHAGE2 based lentivirus vector where the 3’ LTR had been repaired (plasmid map 3274_pH2rU3_ForInd_Pur_Nipah_RBP_CTdel33). Repair of the 3’ LTR enabled us to rescue viruses from cell-stored pseudovirus libraries, a requirement in our system to make genotype-phenotype linked pseudoviruses. This Nipah RBP cytoplasmic tail deletion is based on previous work showing NiV pseudotyped lentivirus titers are greatly increased when most of the cytoplasmic tail is removed (see also <u>fig S3</u>) (*35*, *36*). Rescue of lentiviruses with this vector is dependent on co-transfection of four plasmids (see plasmid maps: 26_HDM_Hgpm227, 27_HDM_tat1b, 28_pRC_CMV_Rev1b, and 3263_HDM_Nipah_F_CTdel22 that contained a 22 amino acid deletion in the cytoplasmic tail but was otherwise unmutated). These plasmids express Nipah F, along with the HIV-1 proteins for Gag/Pol, Tat, and Rev, which are necessary to produce infectious pseudovirus. Genes encoding the different HIV-1 proteins are located on different plasmids and codon optimized to limit the possibility of recombination leading to replication competent HIV. Although our Nipah RBP sequence was downstream of a dox inducible TRE3G promoter, tests showed titers were not improved during rescue when dox was included, suggesting a sufficient amount of RBP is produced from Tat transfection alone. Thus, dox was not used during the rescue of pseudovirus libraries.

### RBP mutant library design and production

We sought to include all possible missense mutations in the ectodomain of the Nipah RBP. The location of the RBP ectodomain was inferred by computational prediction of transmembrane motifs in the RBP amino-acid sequence by DeepTMHMM (*55*). Based on these predictions, the transmembrane domain is located between RBP sites 49-69 (fig S10). To generate the library, we ordered a single mutant library from Twist Biosciences where reference sites 71-602 were mutagenized, and every other position from sites 71-234 included a stop codon. The stop codons were included at a subset of sites to serve as a meaningful negative control, but not so many as to put too many inactive RBPs into the variant libraries. The final QC report from Twist indicated a total of 19 mutants had failed and were not present in the library (https://github.com/dms-vep/Nipah_Malaysia_RBP_DMS/blob/master/data/custom_analyses_data/paper_supplementary_data/twist_QC_report.csv). In order to have biological replicates we made two separate libraries, called LibA and LibB, which were separately barcoded using PCR with a 16 base random barcode. The PCR conditions are as follows. 25 µL of KOD Hot Start Master Mix (ThermoFisher, Cat. No. 71842-4), 1.5 µL of 10 µM each primer (ForInd_AddBC_2, 5’_for_lib_bcing), 5 ng of mutant library template DNA, and 19 µL of water. The thermocycler conditions were:

1. 95°C for 2 min
2. 95°C for 20 seconds,
3. 55.5°C for 20s with ramp rate of −0.5°C/sec,
4. 70°C for 1 minute.
5. Return to step 2 for 9x cycles
6. 12°C hold

The vector was prepared by cutting plasmid 3260_pH2rU3_ForInd_mCherry_T7_CMV_ZsGT2APurR with restriction enzymes XbaI and MluI-HF (NEB, Cat. No. R0145S and R3198S). PCR products and cut vector were run on a 1.5% agarose gel, and bands of the correct size were excised and cleaned up with a NucleoSpin Gel and PCR Clean-up kit (Macherey-Nagel, Cat. No. 740609.5) followed by cleanup with Ampure XP beads (Beckman Coulter, Cat. No. A63881). Barcoded mutant variants were then cloned into the cut vector using the NEBuilder HiFi DNA Assembly Kit (NEB, Cat. No. E2621) with a 1:3 insert to vector molar ratio in a 1 hour reaction. HiFi reactions were purified with Ampure XP beads, and DNA was eluted into molecular grade water. We transformed the purified HiFi reaction into 10-beta electrocompetent cells (NEB, Cat. No. C3020K) with a BioRad MicroPulser Electroporator (Cat. No. 1652100), shocking at 2 kV. Bacteria were then plated on 15 cm LB+amp plates and grown overnight at 37°C. To not bottleneck our library diversity at this stage, we sought to make plasmid libraries with a large number of separate colonies. Total colonies for LibA and LibB were 2×10^6^ and 3×10^6^, respectively. Bacteria colonies were scrapped off the plate and eluted in LB+ampicillin, followed by DNA extraction with a Qiagen HiSpeed Plasmid Maxi Kit (Cat. No. 12662).

### Rescue of RBP pseudovirus library from transfection and creation of cell-stored mutant library

In order to perform DMS with lentiviruses, we needed to be able to link the genotype and phenotype of RBP variants contained in the vector and the protein expressed on the surface of each virion (See <u>fig S1C</u>). Pseudoviruses rescued from transfection will contain a mixed population of RBP proteins on the surface of each virion. Additionally, HIV is pseudodiploid and contains two copies of each RNA genome which necessitates production of pseudoviruses from a single vector sequence in each cell to ensure recombination does not scramble the link between barcodes and specific mutations. Therefore, we used a method (originally described in Dadonaite et al (*33*)) to integrate each pseudovirus as a single copy in a cell, which enables us to generate full genotype-phenotype linked pseudovirus libraries.

The specific method is as follows. 1 µg of the RBP mutant library lentiviral backbone, 250 ng of each lentiviral helper plasmid (26_HDM_Hgpm227, 27_HDM_tat1b, 28_pRC_CMV_Rev1b), and 250 ng of VSV-G (29_HDM_VSV_G) were transfected into 293T cells plated on 6-well plates using BioT transfection reagent (Bioland Scientific, Cat. No. B01-02). After 48 hours, the supernatant was filtered through a 0.45 µm syringe filter (Corning, Cat. No. 431220) and stored at −80°C.

The final pseudovirus variant libraries require a specific number of variants to ensure accurate DMS measurements. If too few variants are present in a library, we would either fail to detect certain mutations, or only have variants associated with a small number of barcodes, which limits measurement accuracy. If too many variants are present, we would have difficulty obtaining high enough pseudovirus titers to ensure all variants are measured during the selection conditions. Therefore, we controlled variant library size by bottlenecking the virus produced from transfections above.

Briefly, an aliquot of virus was thawed and used to infect 293T cells to obtain a titer (transcription units per mL, TU/mL) based on percent positive ZsGreen cells by flow cytometry. Next, this virus was used to infect M3 293T-rtTA cells, which are stable clones expressing reverse tetracycline transactivator (rtTA) and have been selected to maximally express virus from individual proviruses as previously described in Dadonaite et al (*33*). Cells were infected at 0.5% multiplicity of infection (MOI), to ensure each cell contains at most one integrated provirus. Based on estimates for the number of cells present at infection, along with the MOI, we estimated there were ∼60,000 unique variants for each library. The MOI was confirmed by flow cytometry based on the percent of cells that were ZsGreen positive, and cells were subjected to 1 µg/mL puromycin selections for 1 week (changing media, passaging cells, adding fresh puromycin every 48 hours) until only ZsGreen positive cells were observed under a fluorescent microscope. We note that the final number of unique variants present in each library were close to the original 60,000 we estimated above (LibA had 78,450 unique barcodes, LibB had 60,623 variants; fig S4). We ensured the library was not bottlenecked by passaging a minimum of 10 million cells each time. Cells were frozen in the gas phase of liquid nitrogen in aliquots of 10 or 20 million cells / mL.

### Rescue of RBP and F expressing pseudoviruses from cell-stored mutant library

Viral selections require a high titer of pseudoviruses to ensure there is no bottlenecking of the library diversity, which would skew the measurements. To get high enough titers, we rescued pseudoviruses from a large number of cells, followed by concentration to obtain the final library that can be used to perform selections. Specific methods are as follows. To rescue virus from our integrated cell libraries, cells were grown in 5-layer flasks (Corning Falcon 875cm^2^ Rectangular Neck Cell Culture Multi-Flask, Cat. No. 353144) in 150 mL of D10 until 70% confluent. For each 5-layer flask, a transfection mix was prepared as follows. For Nipah pseudovirus, 60 µg of each helper plasmid (26_HDM_Hgpm227, 27_HDM_tat1b, 28_pRC_CMV_Rev1b) were combined with 8 µg of plasmid 3263_HDM_Nipah_F_CTdel22, 9.2 mL of sera free DMEM, and 276 µL of BioT transfection reagent. After incubating at room temperature for 15 minutes, the transfection mix was added to the 5-layer flasks. For VSV pseudovirus, the same procedure as above was used except plasmid 29_HDM_VSV_G replaced the 3263_HDM_Nipah_F plasmid. After 40 hours, the supernatant was filtered through a 0.45um SFCA Nalgene 500mL Rapid-Flow filter unit (Cat. No. 09-740-44B). We concentrated the supernatant by adding Lenti-X Concentrator (Takara, Cat. No. 631232) at a ratio of 1:3, incubating at 4°C for 4 hours, followed by spinning for 45 minutes at 1500 rcf at 4°C. The supernatant was discarded, and viral pellets were resuspended in F-12K media. Following concentration, the titer of concentrated library virus was ∼1*10^6^ TU/mL, and frozen in 1 mL aliquots at −80°C.

### Long-read sequencing to link mutations to barcodes

To link RBP variants with barcodes, we sequenced the entire RBP plus a 16-bp random nucleotide barcode at the 5’ end using PacBio long-read sequencing (see <u>fig S1B</u>; protocol originally described in Dadonaite et al (*33*)). The cell-stored libraries were transfected and viruses were rescued as above, except using 29_HDM_VSV_G in place of 3263_HDM_Nipah_F_CTdel22. 293T cells were then infected with ∼5-10×10^6^ TU of virus. After 12-16 hours, episomal DNA was extracted (see section *Extraction of lentiviral episomal DNA*). 1st round PCR conditions were done in two separate reactions that add a single nucleotide base (either G or C) at the 5’ and 3’ end of the full amplicon to identify strand exchange, so those variants can be filtered downstream. 1st round PCR conditions are as follows. 20 µL of KOD enzyme, 10 µL of DNA prepared from miniprep, 1 µL each of BL_PacBio_5pri_G and BL_PacBio_3Pri_C (both at 10uM) or 1 µL each of BL_PacBio_5pri_C and BL_PacBio_3pri_G, and 8 µL of molecular grade water. The thermocycler conditions were:

1. 95°C for 2 min
2. 95°C for 20 seconds
3. 70°C for 1 second
4. 60°C for 10 seconds with a ramp rate of −0.5°C/sec,
5. 70°C for 1 minute.
6. Return to step 2 for 7x cycles
7. 12°C hold

The PCR products of the two separate reactions for each library were pooled and cleaned up with Ampure XP beads at a 0.8:1 bead to DNA ratio. The conditions for the 2nd round PCR are as follows. 25 µL of KOD enzyme, 2 µL of each primer at 10 µM (5’_PB_Rnd2 and 3’_PB_Rnd2), and 21 µL of cleaned round 1 product. The thermocycler conditions were:

1. 95°C for 2 min
2. 95°C for 20 seconds
3. 70°C for 1 second
4. 60°C for 10 seconds with a ramp rate of −0.5°C/sec,
5. 70°C for 1 minute.
6. Return to step 2 for 10x cycles
7. 12°C hold

PCR products were then cleaned with Ampure XP beads at a 0.8:1 ratio, followed by sequencing on a PacBio Sequel IIe.

To analyze the PacBio sequencing data and link barcodes to variants, we used *alignparse* (https://github.com/jbloomlab/alignparse) (*56*). Sequencing reads were first filtered by error rate (*max_ccs_error_rate <= 1e-4*, which removed approx. 20% of reads for LibA and B). CCSs that did not contain a barcode (∼2% of reads), or that were the result of strand exchange (∼<1%), as identified by the information contained within the C or G nucleotides added during PCR, were also filtered. Next, CCSs were aligned to a Nipah RBP vector reference sequence, and consensus sequences were generated that contained at least three CCSs and less than a fraction of 0.2 for minor sub or indel frequencies within each consensus sequence. A barcode-variant lookup table was made based on the these consensus sequences, with an empirical accuracy of 0.8 for LibA and LibB and is available at the link here: (https://github.com/dms-vep/Nipah_Malaysia_RBP_DMS/blob/master/results/variants/codon_variants.csv). Jupyter notebooks are available for the specific PacBio filtering steps (https://dms-vep.org/Nipah_Malaysia_RBP_DMS/notebooks/analyze_pacbio_ccs.html), the building of PacBio consensus sequences (https://dms-vep.org/Nipah_Malaysia_RBP_DMS/notebooks/build_pacbio_consensus.html), and the generation of the barcode-variant tables (https://dms-vep.org/Nipah_Malaysia_RBP_DMS/notebooks/build_codon_variants.html)

### Extraction of lentiviral episomal DNA

In order to recover information about which variants entered cells following each selection condition, we needed to be able to sequence the barcode sequence from each infecting pseudovirus. Although it might be possible to sequence integrated provirus following infection, the high amount of gDNA that would be co-extracted from cells poses a difficulty due to PCR efficiency. Therefore, we used a method (as originally described in Dadonaite et al (*33*)) to specifically extract episomal lentiviral DNA following cell entry and reverse transcription but prior to integration, which ensures a higher fraction of extracted DNA is of lentiviral origin, rather than host cell. Prior to integration, episomal lentiviral dsDNA will be present in the nucleus as low molecular weight DNA, which can be selectively extracted from cells using a plasmid DNA extraction kit. The specific details are as follows: 5*10^5^ CHO-bEFNB2 or CHO-bEFNB3 cells were plated in 6-well plates in 2 mL of F12-K media. Pseudoviruses were added (refer to specific sections on each type of selection for more information). For all selection conditions, 12-16 hours after infection, cells were trypsinized and spun down at 300 rcf for 4 minutes. We then extracted episomal DNA using a Qiagen QIAprep Spin Miniprep Kit (Cat. No. 27106), following manufacturer’s directions and eluted into 30 µL of Buffer EB.

### Illumina barcode sequencing

To amplify lentivirus genome barcodes, we added 22 µL of extracted episomal DNA (described in the section *Extraction of lentiviral episomal DNA*) to a reaction containing 25 µL KOD, and 1.5 µL each of 10 µM Illumina_Rnd1_For and Illumina_Rnd1_Rev primers. The thermocycler conditions were:

1. 95°C for 2 min
2. 95°C for 20 seconds
3. 70°C for 1 second
4. 58°C for 10 seconds with a ramp rate of −0.5°C/sec,
5. 70°C for 20 seconds.
6. Return to step 2 for 27x cycles
7. 12°C hold

First round products were then cleaned with Ampure XP beads at a ratio of 1:1. 2 µL of cleaned 1st round product diluted to 5 ng/µL was then mixed with 25 µL KOD, 19 µL of molecular grade water, and 2 µL each of 10 µM Illumina_Rnd2_Fwd and a unique index contained with the Illumina_Rnd2_Index_Rev primer. Thermocycler conditions were almost identical to the first round, except using 20 total cycles.

DNA concentration was determined with a Qubit 4 Fluorometer (ThermoFisher, Cat. No. Q33238), and PCR products were pooled in equal DNA amounts, followed by running on a 2% Agarose Gel. The correct band size (283bp) was clipped on a blue light gel dock, followed by extraction with a NucleoSpin Gel and PCR Clean-up kit. The final DNA library was then cleaned up with Ampure XP beads at a ratio of 1:1, and the DNA concentration was determined on a Qubit 4. The final library was diluted to 4 nM, and single read sequencing was done with a NextSeq P1 or P2 kit for 50 cycles. To account for sequencing errors and to ensure each variant was sequenced multiple times (to obtain accurate frequency estimates of each barcode), we oversequenced each selection, and recovered ∼10-85 million reads total. Thus, each barcode in a selection was sequenced on average 90-600 times, depending on the exact read depth. Details on the number of Illumina reads mapping to each selection, and the average number of reads per barcode are given in this notebook (https://dms-vep.org/Nipah_Malaysia_RBP_DMS/notebooks/analyze_variant_counts.html).

### Validations of DMS data using single mutant Nipah pseudoviruses

To validate our DMS measurements, we made single RBP mutant pseudoviruses and compared their entry or neutralization to unmutated Nipah RBP. For RBP entry, we made RBP mutants C162L, F168A, Y389T, P488S, Q492L, Q530F, Q530E, and Q530L. For receptor neutralization validations, we made mutants H333Q, Q492R, V507I, Q530F, S553W, and D555K. For antibody neutralization, we made mutants L184H, P185D, Q450S, D468V, I517K, W519T, and I520T.

Specifically, we introduced single amino-acid mutations using PCR with partially overlapping primers that contained the desired mutations. Primers were designed with the online NEBaseChanger software. For PCR, 10 ng of plasmid 3336_HDM_Nipah_RBP_CTdel33 was added to two reactions containing 25 µL KOD, and 1.5 µL each of 10 µM For_primer and Rev_mutagenesis_primer or Rev_primer, Forward_mutagenesis_primer. The thermocycler conditions were:

1. 95°C for 2 min
2. 95°C for 20 seconds
3. 58°C for 10 seconds
4. 70°C for 30 seconds.
5. Return to step 2 for 24x cycles
6. 12°C hold

Following PCR, the two amplicons were cloned into plasmid 27_HDM_tat1b after cutting with NotI and HindIII, with the NEBuilder Hifi DNA Assembly Kit incubating at 50°C for 1 hour, and transformed into Stellar competent cells (Takara, Cat. No. 636763). For cell entry validation mutants, three separate plasmids were isolated; for receptor binding or antibody neutralization validation only one plasmid was isolated per mutant. All mutants were confirmed by whole plasmid sequencing by Primordium. To make virus, 293T cells were plated on 6-well plates in 2mL of D10 media. 16-24 hours later, 1ug of plasmid 2727_pHAGE6-wtCMV-Luc2-BrCr1-ZsGreen-W-1247, 500ng of 26_HDM_Hgpm2, and 250ng each of 3263_HDM_Nipah_F_CTdel22, and either the unmutated RBP expression plasmid 3336_HDM_Nipah_RBP_CTdel33, or a mutant expression plasmid. An example of a mutant expression plasmid is 3922_HDM_Nipah_RBPctDel_P185D. After 48 hours, the supernatant was passed through a 0.45 uM filter.

For cell entry validations, three separate plasmid preps of each RBP variant were used to generate three separate virus stocks. Viruses were added to either CHO-bEFNB2 or CHO-bEFNB3 cells plated a day earlier in poly-L-lysine coated, black walled, 96-well plates (Greiner, Cat. No. 655930). For each plasmid prep, two replicates were run on the same plate at three different dilutions to ensure we were in the correct dynamic range. Each plate also contained a cell only and virus only condition to correct for any background luciferase signal. 48 hours later, 170 µL of supernatant was removed from each well, followed by the addition of 30 µL of Bright-Glo Luciferase Assay System (Promega, E2610). Luciferase values were immediately read on a Tecan Infinite M1000 plate reader. The relative light units (RLU) per µL were calculated separately for each mutant. The data for validation assays for CHO-bEFNB2 cells are here (https://github.com/dms-vep/Nipah_Malaysia_RBP_DMS/blob/master/data/custom_analyses_data/experimental_data/functional_validations_EFNB2.csv) and for CHO-bEFNB3 cells here (https://github.com/dms-vep/Nipah_Malaysia_RBP_DMS/blob/master/data/custom_analyses_data/experimental_data/functional_validations_EFNB3.csv).

To validate DMS measurements for antibody and receptor neutralization, the same strategy was used as above, except luciferase expressing pseudoviruses were incubated with antibody or soluble receptor for one hour prior to adding to cells. For antibody validations, we used a starting concentration of 10 ug/mL of nAH1.3 and did eight three-fold dilutions. A no-antibody or no-receptor control were run in duplicate. Fraction infectivity was calculated by subtracting background readings and comparing the amount of signal in each well with the average luciferase reading from two wells with no antibody or no receptor. Fraction infectivity values for the antibody validations are here (https://github.com/dms-vep/Nipah_Malaysia_RBP_DMS/blob/master/data/custom_analyses_data/experimental_data/nAH1_3_mab_validation_neuts.csv), and for receptor validations here (https://github.com/dms-vep/Nipah_Malaysia_RBP_DMS/blob/master/data/custom_analyses_data/experimental_data/binding_single_mutant_validations.csv). IC_50_ values were estimated by fitting neutralization curves using the package *neutcurve* (https://github.com/jbloomlab/neutcurve) (*57*).

### Production of a VSV-neutralization standard virus

To estimate absolute neutralization across different conditions, we used pseudoviruses expressing VSV-G that were not neutralized by ephrin receptors or anti-RBP antibodies, and spiked them into the receptor binding and antibody selections to represent ∼1% of reads in the no-antibody or no-receptor control selections (see below). The VSV-G neutralization standard viruses were produced as described in Dadonaite et al (*33*). Briefly, a sequence encoding mCherry that was associated with eight known 16bp barcodes was cloned into a lentiviral vector (plasmid 3260_pH2rU3_ForInd_mCherry_T7_CMV_ZsGT2APurR). These viruses were rescued by transfection with plasmid expressing VSV-G and the other helper plasmids in 293T cells, followed by transduction of M3 293T-rtTA cells at low MOI, to ensure no more than one vector was integrated per cell. Pseudoviruses were then rescued as described above, and titered on 293T cells to be able to spike in a known amount relative to the RBP/F pseudovirus libraries.

### Selections to determine effects of mutations on cell entry

To determine the effects of mutations on cell entry, we used a strategy where barcode frequencies between VSV-G pseudotyped lentiviral mutants are compared with barcodes for pseudoviruses with Nipah RBP and unmutated F (see also fig <u>S7A</u>). Because VSV-G efficiently infects cells irrespective of which RBP variant is on the virion surface, it is used as a ‘control’ for determining the baseline composition of the mutant library. RBP mutants will have variable ability to enter cells and can thus be compared with the baseline composition.

Specifically, we plated 7.5*10^5^ CHO-bEFNB2 or CHO-bEFNB3 in 2mL of F12-K media in individual wells of a 6-well plate. The next day, ∼5*10^6^ VSV-G pseudotyped viruses were added to cells, or ∼1*10^6^ Nipah RBP/F pseudoviruses. 12-16 hours later, episomal DNA was extracted from cells (see section *Extraction of lentiviral episomal DNA*), followed by Illumina sequencing (see section *Illumina barcode sequencing*).

We performed at least two independent cell entry selections from each library, although in some cases we included more to improve confidence of DMS measurements. For estimating cell entry in CHO-bEFNB2 cells, we performed six LibA and two LibB selections independently. For estimating cell entry in CHO-bEFNB3 cells, we performed three LibA and four LibB selections.

To analyze the Illumina sequencing data, we used the package *dms_variants* (https://github.com/jbloomlab/dms_variants) to parse the read data and estimate functional scores associated with each variant. Barcodes were aligned to the codon-variant table previously produced from PacBio long-read sequencing. Raw barcode counts were converted to functional scores as previously described in Dadonaite et al. (*33*). Raw barcode counts can be found here (https://github.com/dms-vep/Nipah_Malaysia_RBP_DMS/tree/master/results/barcode_counts). Because our libraries contained some multi-mutants, we deconvolved the effects of individual mutations on entry with a global epistasis model (*40*) implemented in the *multidms* package (https://github.com/matsengrp/multidms). The final entry score we report is taken from the average effect across replicates and libraries.

### Selections to determine effects of mutations on receptor binding

To determine the effects of mutations on receptor binding, we used a strategy where pseudoviruses were incubated with varying amounts of soluble receptor, followed by cell infection to recover barcode frequencies for the different conditions (see <u>fig S7B</u>). The underlying reasoning is that pseudovirus neutralization by soluble receptors should be proportional to its binding. Mutations which increase receptor binding will be neutralized at a lower concentration than unmutated, while mutations which decrease binding will be neutralized at a higher concentration.

Specifically, we plated either 7.5*10^5^ CHO-bEFNB2 or CHO-bEFNB3 in 2mL of F12-K media in individual wells of a 6-well plate. The next day, the mutant pseudovirus library was mixed with 1% of a VSV-G pseudovirus ‘neutralization standard’ which contains eight different known barcodes (see section *Production of a VSV-neutralization standard virus*). Because VSV-G is not neutralized by soluble bEFNB2 or bEFNB3, its proportion of reads in each condition will vary depending on neutralization of RBP pseudovirus, which allows us to standardize reads across conditions. Next, ∼1*10^6^ TU of Nipah RBP/F pseudoviruses were used to either infect cells (control condition) or incubated with varying amounts of soluble monomeric bEFNB2 or dimeric bEFNB3 for one hour. Based on previously determined IC_50_ ranges (Fig 3A), we sought to use concentrations spanning an IC_5_ to IC_95_. Following incubation, pseudoviruses were added to cells (viruses from monomeric bEFNB2 selections infected CHO-bEFNB2 cells, viruses from dimeric bEFNB3 selections infected CHO-bEFNB3 cells). 12-16 hours later, DNA was extracted (see section *Extraction of lentiviral episomal DNA*), followed by Illumina sequencing (see section *Illumina barcode sequencing*).

To analyze the sequencing data, we parsed the Illumina reads as in the previous section. We fit neutralization curves for each selection to estimate the effects of mutations on receptor neutralization as implemented in the package *polyclonal* (https://jbloomlab.github.io/polyclonal/) (*58*). At a minimum, we did at least one selection with LibA and LibB, and used the mean effects between these independent replicates in all analyses.

### Selections to determine effects of mutations on antibody neutralization

To determine the effect of RBP mutations on antibody neutralization, we used the same general strategy as above, except substituting antibodies in place of the soluble receptor proteins. All antibody selections were followed by infection of CHO-bEFNB3 cells. Based on previous IC_50_ values for each antibody (<u>fig S25</u>), pseudoviruses were incubated with antibody concentrations corresponding to IC_50_, IC_90_, and IC_99_. Neutralization curves were fit as in the previous section. At a minimum, we did at least one selection with LibA, and one with LibB, and used the mean effects between these independent replicates in all analyses.

### Data filtering

Raw data were filtered to produce all heatmaps and figures. Briefly, we required mutations to be present in multiple barcodes (*times_seen > 2*). We also filtered out mutations with a high standard deviation between replicates (*max_std*). Finally, we required mutations to be observed in the majority of individual selections (*frac_models > 0.5*). These steps remove mutations that have a low number of observations, with high standard deviation between selections, or which were only observed in a subset of individual selections. Notebook for filtering data is here (https://dms-vep.org/Nipah_Malaysia_RBP_DMS/notebooks/filter_data.html), and contains more information about the specific parameters that were used.

### Validation of RBP binding measurements by biolayer interferometry

To validate our DMS measurements for receptor binding, we expressed monomeric RBP mutants and tested their binding affinity to dimeric bEFNB2 or bEFNB3 by biolayer interferometry (BLI). Specific details are included below.

The Nipah RBP head domain construct contains residues 176-602 from Nipah RBP, Mu phosphatase signal peptide, and a N-terminal GSGGGS followed by a 6x Histidine tag in the pOPING expression vector. The mutations V244W, L305W, Q492L, V507I, Q530F, S553W, D555Y, Q559R, and I588V were subcloned into the same NiV RBP head construct described above in the pTwist CMV expression vector. The bEFNB2 and bEFNB3 hFc constructs contain residues 28-224 for ephrin-B2 and 29 to 226 for ephrin-B3; N-terminal MPMGSLQPLATLYLLGMLVASVLA signal peptide; and C-terminal thrombin cleavage site, IEGRMD linker followed by residues 100 to 330 of human IgG1 in the pTwist CMV expression vector.

Nipah RBP heads were expressed in Expi293 cells (Thermo) at 37°C and 8% CO_2_. Cells were transfected using Expifectamine293 transfection kit (Thermo) following the manufacturer’s protocol. Four days after transfection, Expi293 cell supernatant was clarified by centrifugation at 4,121 x *g* for 30 minutes, supplemented with 25 mM phosphate pH 8.0, 300 mM NaCl. Supernatant was then bound to Ni Excel resin (Cytiva) previously equilibrated in 25 mM phosphate pH 8.0, 300 mM NaCl. Nickel resins were washed with 30 mL of 25 mM phosphate pH 8.0, 300 mM NaCl, and 20 mM imidazole. Protein was eluted using 25 mM phosphate pH 8.0, 300 mM NaCl, and 500mM imidazole prior to being buffer exchanged to 50 mM Tris-HCl pH 8.0, 150 mM NaCl using a centrifugal filter device with a MWCO of 10 kDa. Protein was then flash frozen and stored at −80°C until use. Proteins were run over a Superdex200 increase 10/300 size-exclusion column (Cytiva) equilibrated in 50 mM Tris pH 8.0 and 150 mM NaCl. Fractions containing monodisperse protein were then used for biolayer interferometry.

Biolayer interferometry (BLI) was performed on an Octet Red96 (Sartorius) and the Octet Data acquisition. Dimeric *P. alecto* Ephrin B2-hFc and B3-hFc were diluted to 10 µg/mL in 10x Octet kinetics buffer (Sartorius). Ephrins were then loaded onto hydrated anti-Human-Fc capture (AHC) biosensors to a 1nm shift, equilibrated in 10x Octet kinetics buffer for 60 seconds, and dipped into monomeric NiV G head at 50 nM for ephrin B2 and at 100 nm for ephrin B3. The association phase was run for 300s. Dissociation was observed by dipping biosensors in a 10x Octet kinetics buffer for 300s. BLI measurement was performed at 30°C and shaking at 1,000 rpm. Association phases were aligned to 0 seconds and 0 shift in Octet Data Analysis HT software and the processed results were exported. Sensorgrams were plotted in GraphPad Prism10. Area under the curve and max response was calculated in GraphPad Prism10 analysis.

### *Pteropus alecto* Ephrin B2-hFc and B3-hFc production

In order to test the neutralizing potency of different bat ephrins on unmutated RBP/F pseudovirus, and to perform selections on variant libraries with soluble bat ephrins, we produced soluble ephrin proteins as follows.

Dimeric *Pteropus alecto* Ephrin B2-hFc and B3-hFc were expressed in Expi293 cells (Thermo) at 37°C and 8% CO_2_. Cells were transfected using Expifectamine293 transfection kit (Thermo) following the manufacturer’s protocol. Four days after transfection, Expi293 cell supernatant was clarified by centrifugation at 4,121 x *g* for 30 minutes, supplemented with 20 mM phosphate pH 8.0, 100 mM NaCl. Supernatant was then bound to HiTrap Protein A column (Cytiva) previously equilibrated in 20 mM phosphate pH 8.0, and 100 mM NaCl. Columns were washed with 30 mL of 20 mM phosphate pH 8.0, 100 mM NaCl. Protein was eluted using 8 mL of 100 mM citric acid pH 3.0 into 2 mL of 1M Tris 9.0 pH. Proteins were concentrated using a centrifugal filter device with a MWCO of 30 kDa and run over a Superdex200 increase 10/300 size-exclusion column (Cytiva) equilibrated in 50 mM Tris pH 8.0 and 150mM NaCl. Fractions containing monodisperse dimeric protein were then flash frozen and stored at −80°C until use.

### Monomeric *Pteropus alecto* Ephrin-B2 and -B3 purification

The monomeric P. alecto EFNB2 and EFNB3 constructs were codon-optimized for a mammalian cell expression system, synthesized, and cloned into a pTwist CMV vector by Twist Bioscience. These constructs include residues 28 to 224 for *P. alecto* EFNB2 (GenBank accession no. NP_001277099), and residues 29 to 226 for P. alecto EFNB3 (GenBank accession no. NP_001277094), followed by Factor Xa protease site (IEGR), a linker (GSGGGS) and StrepII tag (WSHPQFEK). MPMGSLQPLATLYLLGMLVASVLA was used as the signal peptide for both constructs.

The monomeric *P. alecto* EFNB2 and EFNB3 were expressed in Expi293F cells by transient transfection using the ExpiFectamine 293 Transfection Kit (Thermo Fisher) according to the manufacturer’s protocols. After 7 d in a humidified shaking incubator maintained at 37 °C and 8% CO2, the transfected cells were harvested and cleared of cellular debris by centrifugation for 10 min at 1,000 × g followed by centrifugation for 30 min at 10,000 × g. The supernatants were then subjected to affinity chromatography. The monomeric P. alecto EFNB2 and EFNB3 were purified from clarified supernatants using a 1-mL StrepTrap HP column (Cytiva), buffer-exchanged, concentrated, and flash-frozen in TBS (pH 8.0, 25 mM Tris, 150 mM NaCl). All columns were equilibrated in TBS (pH 8.0, 25 mM Tris, 150 mM NaCl). The wash buffer used for all columns was TBS. The elution buffer was TBS with 2.5 mM desthiobiotin.

### Logo plots

To make logo plots visualizing antibody escape (i.e. Fig 4A), we used the *dmslogo* package (https://github.com/jbloomlab/dmslogo). To select a subset of sites for visualization, we required mutations to have a max escape score within at least 50% of the max escape score for that antibody, or a summed escape score within at least 75% of the max summed escape site for that antibody.

### Structural analyses

UCSF ChimeraX v1.6.1 (*59*) was used for all structural analyses and figures. PDB accession IDs for specific structures are given in each figure legend. In some cases, superimposition of structures was required (ex. Fig 2A, <u>fig S25</u>), which was done using the *matchmaker* method implemented in ChimeraX using default parameters. To map mutational effects onto the structure, we calculated the site-averaged effects using this notebook (https://dms-vep.org/Nipah_Malaysia_RBP_DMS/notebooks/mapping_site_level.html), which were converted to a .defattr file format required by ChimeraX. Sites were colored using the *color byattr* command function in ChimeraX with the appropriate color scale. To estimate atomic distances between Nipah RBP residues and receptors or antibodies, we used this notebook (https://dms-vep.org/Nipah_Malaysia_RBP_DMS/notebooks/receptor_distance.html). Receptor and antibody contact sites were calculated by finding all RBP residues within 4 angstroms.

### Sequence analysis

To obtain evolution and diversity information regarding Nipah viruses (for example <u>fig S2</u>, Fig 4C), we downloaded all publicly available Nipah whole-genome nucleotide sequences from GenBank (as of January-3-2024; GenBank accessions found here https://github.com/dms-vep/Nipah_Malaysia_RBP_DMS/blob/master/data/custom_analyses_data/alignments/phylo/nipah_whole_genome_genbank_accession_ids.txt). Sequences were aligned with MAFFT v7.520 (*60*), and a maximum-likelihood phylogenetic tree was inferred with IQ-Tree v2.2.2.6 (*61*) with the TIM2 substitution model. The phylogeny was then visualized using the package *baltic 0.2.2* (https://github.com/evogytis/baltic) implemented in this notebook (https://dms-vep.org/Nipah_Malaysia_RBP_DMS/notebooks/make_nipah_phylogeny_baltic.html).

To find all RBP amino-acid mutations in these sequences, we trimmed the whole-genome sequences to just RBP and translated them to amino-acid sequences in Geneious Prime 2023.0.4. Alignment can be found here (https://github.com/dms-vep/Nipah_Malaysia_RBP_DMS/blob/master/data/custom_analyses_data/alignments/Nipah_RBP_AA_align.fasta). All amino-acid mutations relative to the unmutated reference sequence were found using this notebook (https://dms-vep.org/Nipah_Malaysia_RBP_DMS/notebooks/henipavirus_conservation.html).

To make an amino-acid alignment of human and bat (*Pteropus alecto*) EFNB2 and EFNB3 sequences (<u>fig S6</u>), we downloaded reference sequences from GenBank and aligned them using MAFFT v7.520. Amino-acid alignment can be found here (https://github.com/dms-vep/Nipah_Malaysia_RBP_DMS/blob/master/data/custom_analyses_data/alignments/ephrin_E2_E3_sequences.fasta). The alignment was visualized using the ENDscript 2 online server (*62*)

### Figures

To make publication quality images, figures were generated with Vega-Altair v5.1.2 (https://altair-viz.github.io/) or Prism v9.4.1. Font sizes and figure arrangements were adjusted in Adobe Illustrator v27.0. Tables were made using the R package *flextable* (https://cran.r-project.org/web/packages/flextable/index.html).

## Notes

https://github.com/dms-vep/Nipah_Malaysia_RBP_DMS

