## Supplemental Figures for "Functional and antigenic landscape of the Nipah virus receptor binding protein"

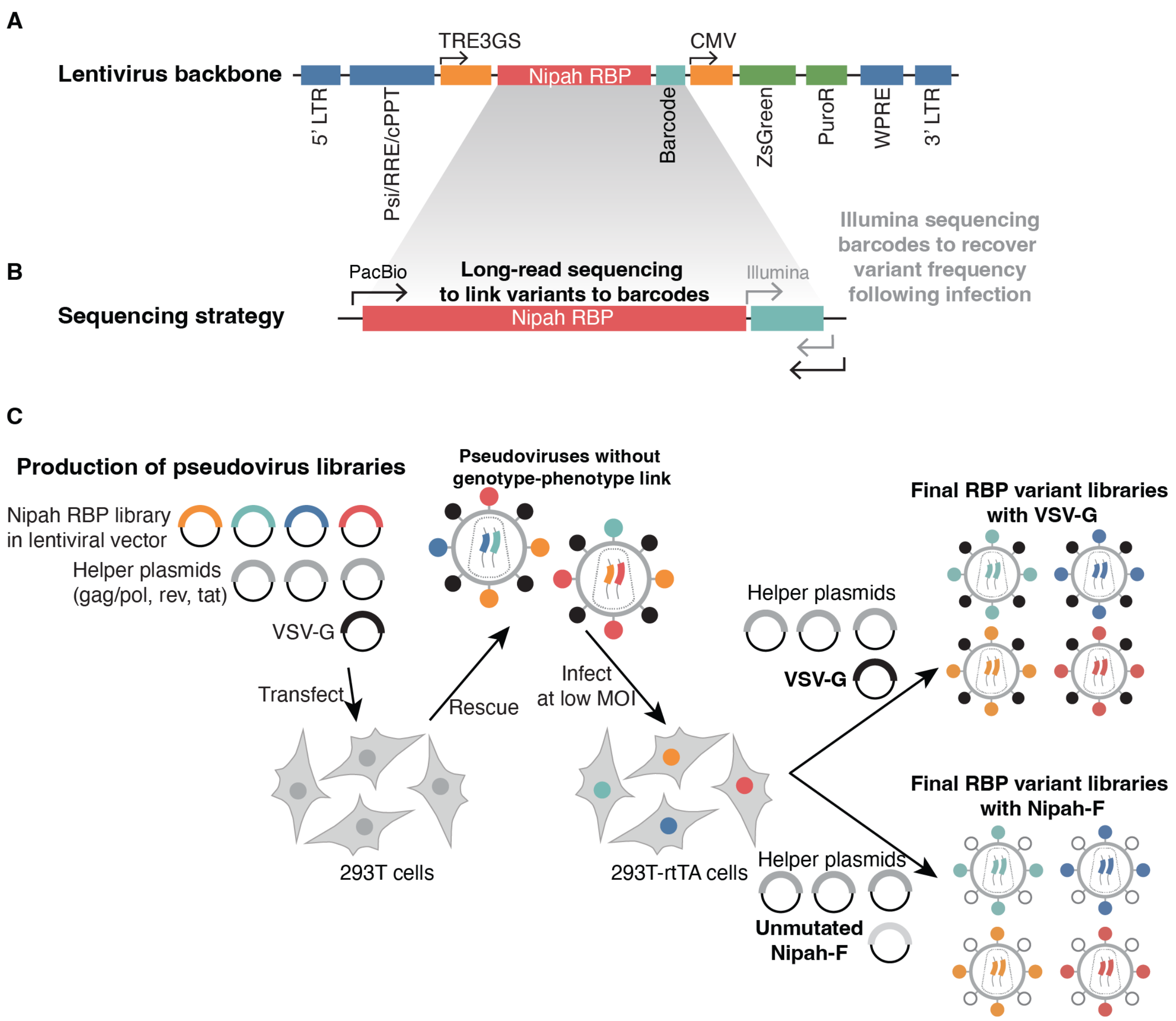


**Supplemental Figure 1. Graphical overview of the lentiviral backbone, sequencing strategy and production of lentiviral libraries pseudotyped with Nipah RBP and unmutated Nipah F**. A) Lentiviral backbone. Mutagenized RBP variants followed by unique 16-nucleotide barcodes were cloned into a lentiviral vector downstream of an inducible TRE3GS promoter. ZsGreen and puromycin resistance genes (PuroR) are downstream of a CMV promoter. The vector also contains essential lentiviral motifs, including the 5' and full-length 3' long-terminal repeats (LTR), the psi packaging element (Psi), rev response element (RRE), and central polypurine tract (cPPT). The Woodchuck hepatitis virus post-transcriptional regulatory element (WPRE) is included for increased transgene expression. B) Sequencing of RBP variants and barcodes. To link variants in RBP with specific barcodes, we PCR-amplified a region that spans the whole RBP gene and barcode region, and then used long-read PacBio sequencing to determine which RBP mutant was linked to each barcode. For DMS experiments, we could then simply PCR amplify and Illumina sequence the short barcode region to identify the full RBP sequence. C) Strategy to make genotype-phenotype linked pseudoviruses. A barcoded library of codon-optimized RBP variants based on the Nipah Malaysia strain was cloned into a HIV-based lentiviral vector (shown in A), and transfected into 293T cells with additional lentiviral helper plasmids (encoding gag/pol, rev, tat) and a plasmid encoding the envelope protein from the vesicular stomatitis virus (VSV-G), which has broad tropism. Pseudoviruses rescued from transfections are not suitable for DMS as they lack a genotype-phenotype link, since each pseudodiploid virion contains two genomes and different RBP variants on the surface. To establish a link between the RBPs on the virion surface and the genotype, we infected 293T-reverse tetracycline-controlled transactivator (rtTA) cells at a low multiplicity of infection (MOI $< 0.01$) to ensure a single integration per cell. Cells containing an integrated provirus are selected with puromycin, creating a cell-stored library of RBP mutants in a lentiviral backbone. Re-transfection of helper plasmids into cell-stored libraries plus either VSV-G or unmutated Nipah-F generates virions that have a single RBP protein variant on their surface and encode an identifying barcode in their genome.

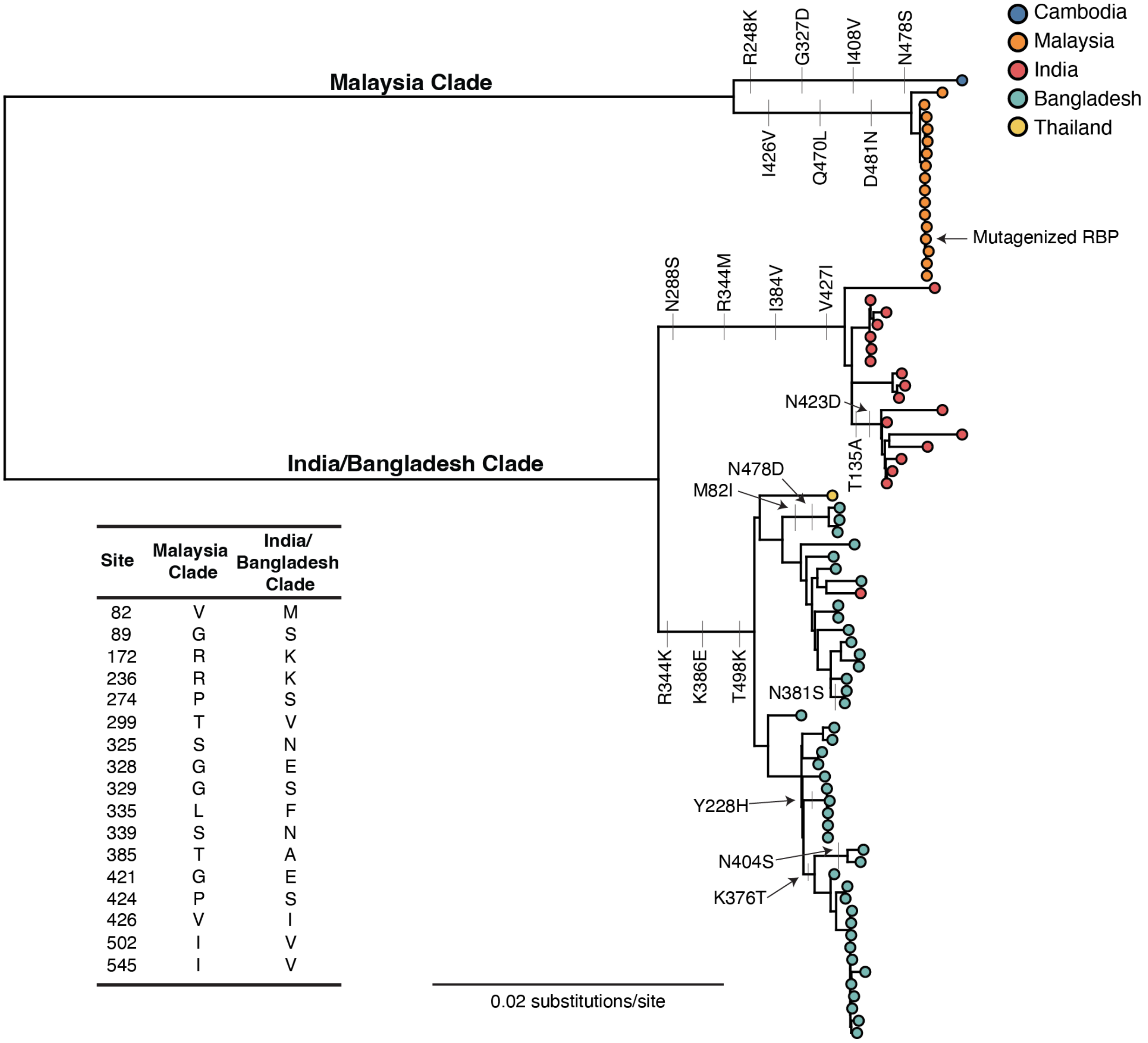


| Site | Malaysia Clade | India/ Bangladesh Clade |
|---|---|---|
| 82 | V | M |
| 89 | G | S |
| 172 | R | K |
| 236 | R | K |
| 274 | P | S |
| 299 | T | V |
| 325 | S | N |
| 328 | G | E |
| 329 | G | S |
| 335 | L | F |
| 339 | S | N |
| 385 | T | A |
| 421 | G | E |
| 424 | P | S |
| 426 | V | I |
| 502 | I | V |
| 545 | I | V |

**Supplemental Figure 2. Evolution and diversity of the Nipah virus RBP**. A maximum likelihood phylogeny was inferred from all publicly available full-length Nipah virus nucleotide sequences downloaded from Genbank (on Jan-3-2024). The tree was inferred with IQ-Tree v2.2.2.6 using a TIM2 substitution model. Scale bar shows the number of nucleotide substitutions per site. Tips are colored by country of origin. We label all RBP amino-acid mutations found in at least two sequences, annotating them directly on the tree or in a separate table when they differ between the two main clades. For the unique Cambodian sequence set apart by a long branch (top taxa on tree), we label all its unique mutations. The Malaysia strain used in our DMS study is indicated with an arrow and text.

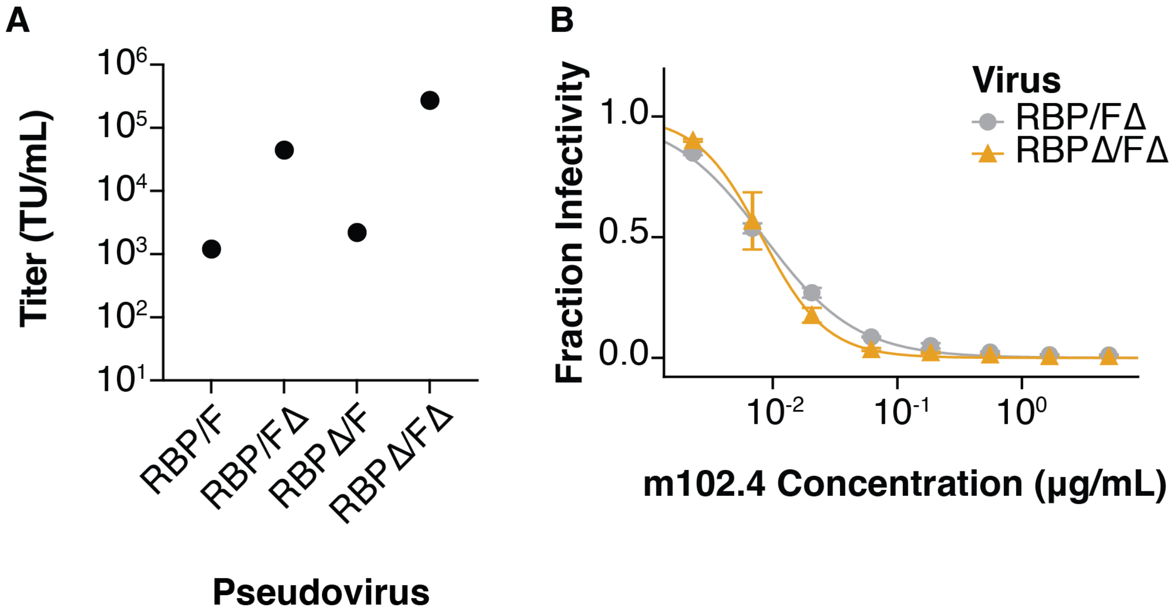


**Supplemental Figure 3. Truncating the cytoplasmic tails of the Nipah virus RBP and F proteins improves titers of pseudotyped lentiviral particles without affecting antigenicity**. Based on prior research on pseudotyping lentiviruses with Nipah RBP/F, we removed 32 and 22 amino acids from the cytoplasmic tails of RBP and F proteins respectively. This modification improved pseudovirus titers, so we utilized the glycoproteins with truncated cytoplasmic tail in all subsequent experiments. A) Titers of unmutated Nipah RBP/F pseudoviruses titered in HEK293T cells with or without cytoplasmic tail truncations (indicated by Δ) in RBP and/or F. B) Neutralization by the anti-RBP monoclonal antibody m102.4 of pseudovirus with full-length RBP compared to RBP with a cytoplasmic tail truncation. The neutralization curves are indistinguishable, suggesting the cytoplasmic tail deletion does not affect antigenicity.

**A**

| Library | Unique barcodes | % mutations present |
|---|---|---|
| LibA | 78,450 | 99.48 |
| LibB | 60,623 | 99.72 |

**B**

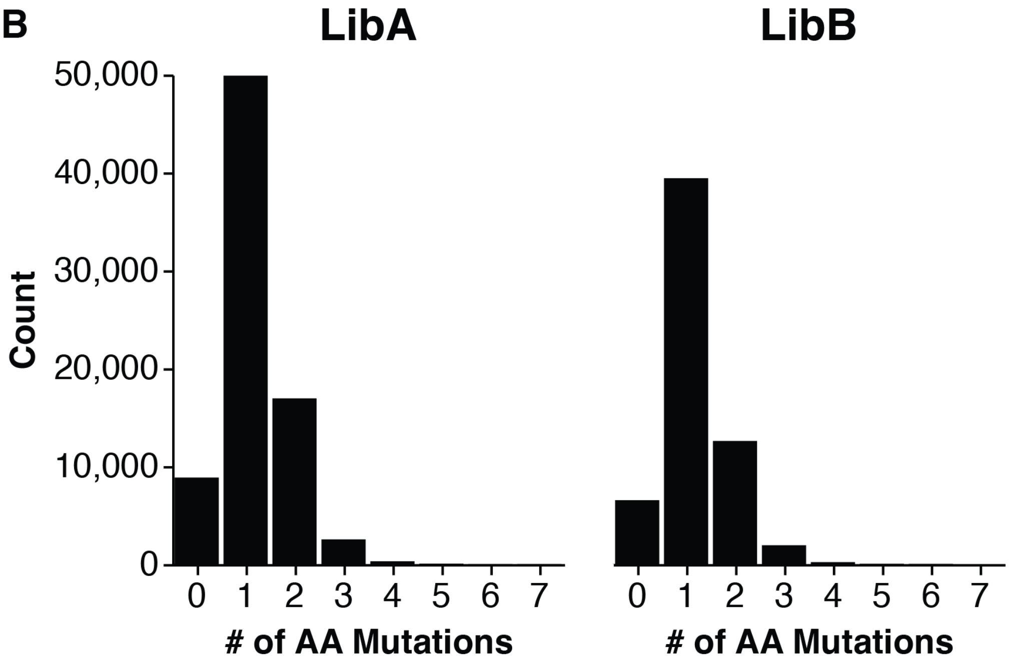


**Supplemental Figure 4. Statistics of RBP mutant library composition.** A) Total number of unique barcodes in each library and the number of amino-acid mutations that were present out of the theoretical total. B) Histogram showing the distribution of amino-acid mutations per variant for each library.

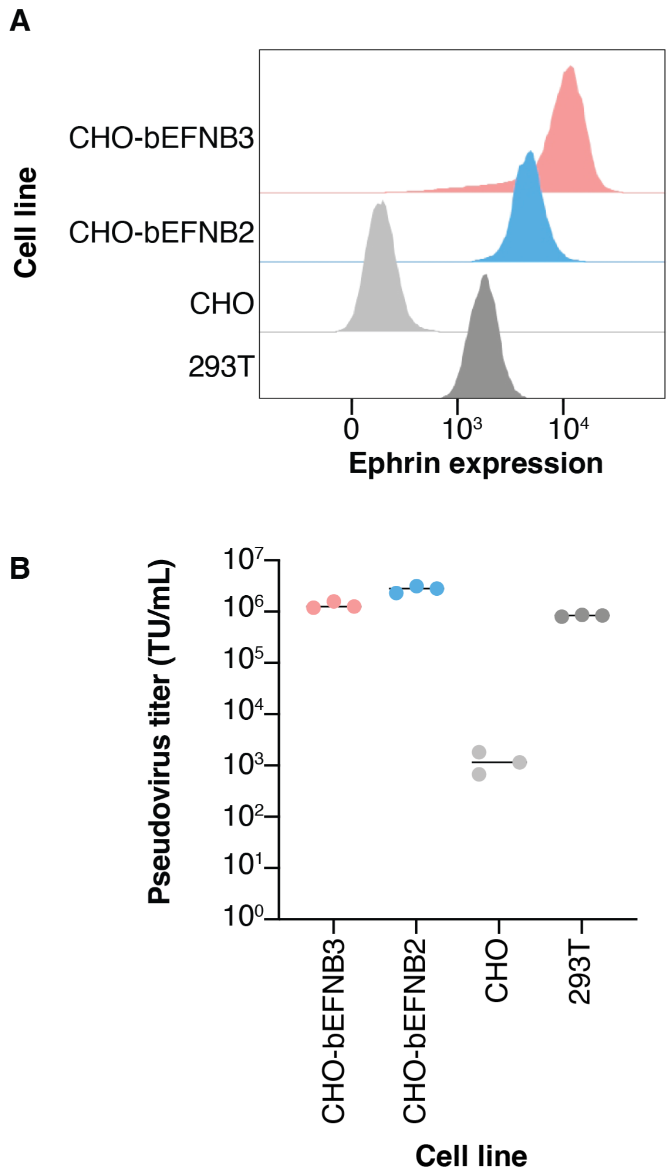


**Supplemental Figure 5. *Pteropus alecto* EFNB2/3 expression levels and susceptibility to infection by unmutated RBP/F-pseudotyped lentiviruses of the stable clones used as target cells in this study.** CHO cells were transduced with a lentiviral vector expressing bat (*Pteropus alecto*) EFNB2 (bEFNB2) or EFNB3 (bEFNB3) downstream of an EF1-alpha promoter, and sorted individual cells were expanded. Clones were selected to have low bEFNB2/bEFNB3 expression (to more closely approximate expression in 293T cells) A) Flow cytometry staining data using EphB3-Fc to determine relative expression of ephrin. EphB3 is a natural ligand of all ephrin-B proteins. Secondary staining was done with a goat anti-human IgG Fc with FITC. B) Titers obtained in different cell lines of lentivirus pseudotyped with unmutated Nipah ΔRBP and ΔF produced by transfection of 293T cells (performed in triplicate).

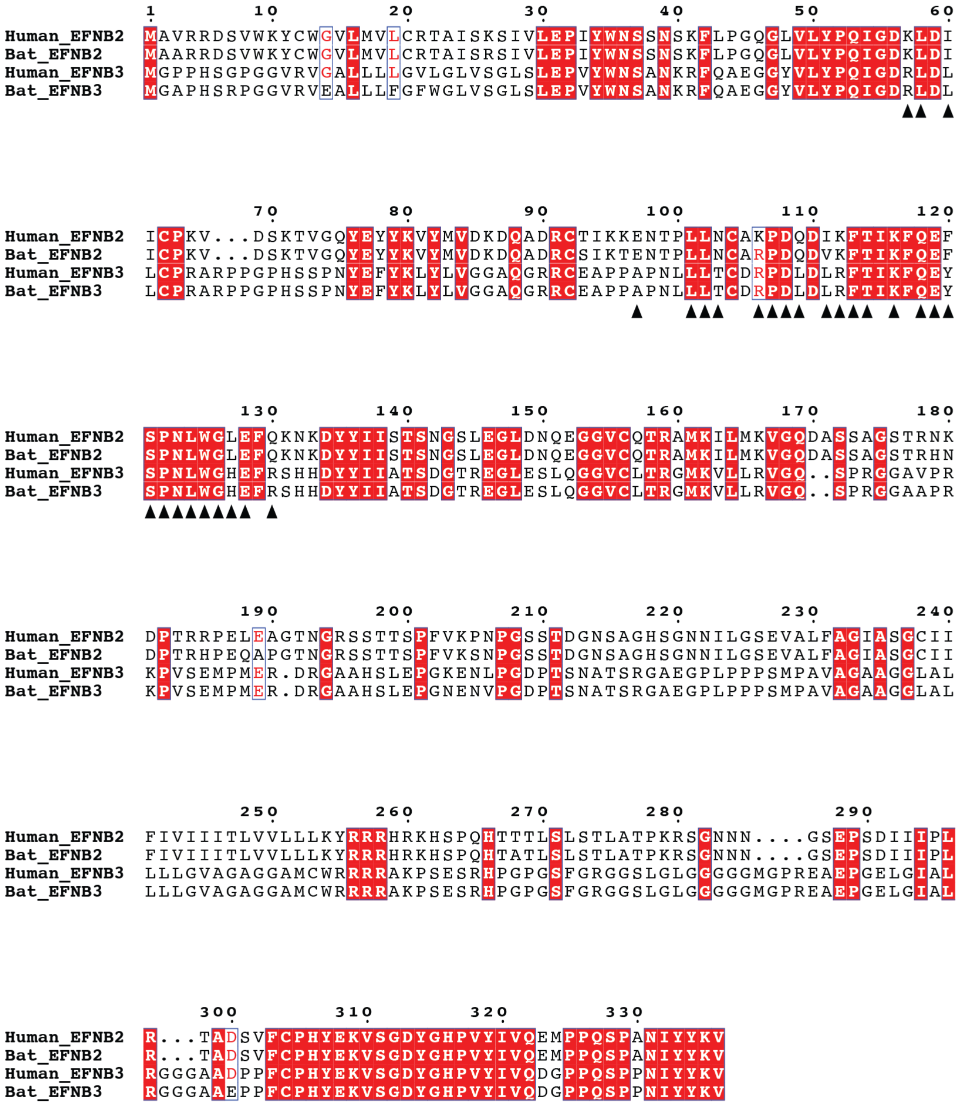


**Supplemental Figure 6. Amino-acid alignment of human and bat (*Pteropus alecto*) EFNB2 and EFNB3 amino-acid sequences**. Ephrin sites that contact RBP residues, defined as less than four angstroms apart (calculated from a PDB structure of RBP in complex with EFNB2 (PDB: 2VSM)) are indicated below the sequence alignment with black triangles. Position numbering is shown above sequences and is based on the EFNB3 sequences. Human and bat EFNB2 contact residues differ at sites 106 and 111. Human and bat EFNB3 contact residues are identical. Sites conserved between all sequences are in red boxes. Sites that differ in only one of the sequences are shown in boxes with red letters. Figure was generated using the ENDscript 2 online server (*62*).

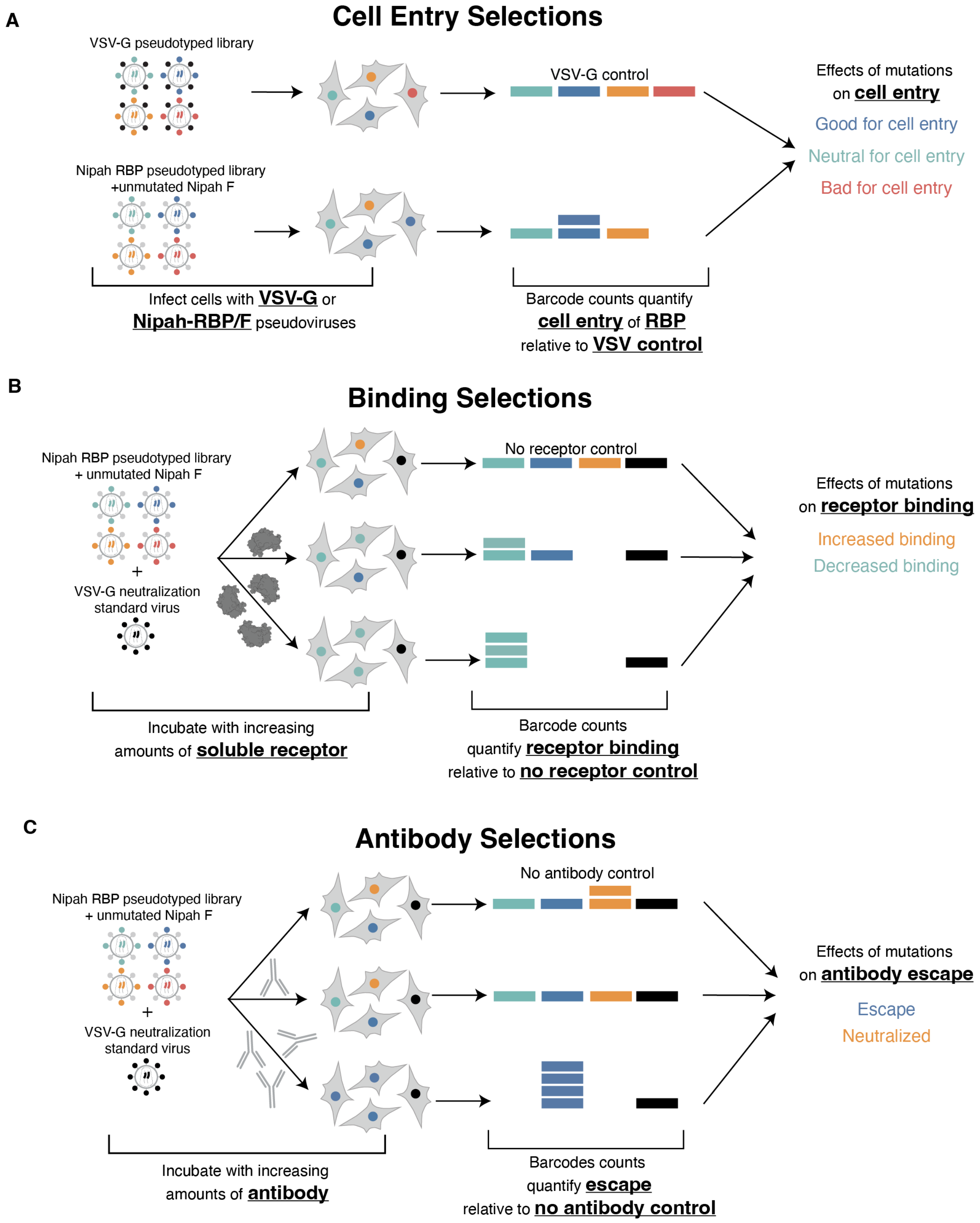

A
Cell Entry Selections
VSV-G pseudotyped library
VSV-G control
Nipah RBP pseudotyped library
+unmutated Nipah F
Infect cells with VSV-G or
Nipah-RBP/F pseudoviruses
Barcode counts quantify
cell entry of RBP
relative to VSV control
Effects of mutations
on cell entry
Good for cell entry
Neutral for cell entry
Bad for cell entry
B
Binding Selections
Nipah RBP pseudotyped library
+ unmutated Nipah F
+
VSV-G neutralization
standard virus
No receptor control
Incubate with increasing
amounts of soluble receptor
Barcode counts
quantify receptor binding
relative to no receptor control
Effects of mutations
on receptor binding
Increased binding
Decreased binding
C
Antibody Selections
Nipah RBP pseudotyped library
+ unmutated Nipah F
+
VSV-G neutralization
standard virus
No antibody control
Incubate with increasing
amounts of antibody
Barcodes counts
quantify escape
relative to no antibody control
Effects of mutations
on antibody escape
Escape
Neutralized

**Supplemental Figure 7. Graphical overview of DMS selections for measuring cell entry, receptor binding, and antibody escape.** A) Cell entry overview. Pseudoviruses with either Nipah-RBP + VSV-G (control condition) or Nipah-RBP + unmutated Nipah-F are used to infect target cells (CHO-bEFNB2 or CHO-bEFNB3). Pseudoviruses with VSV-G on the surface will efficiently infect cells regardless of which RBP variant is expressed on the surface of the virion. 12 hours after infection, unintegrated viral DNA is extracted and barcodes within each lentivirus vector are amplified with PCR and sequenced with Illumina. Reads from the VSV-G control conditions are used to quantify cell entry relative to the Nipah RBP + unmutated F infection condition. B) Receptor binding overview. This approach is based on the fact that inhibition of infection by soluble receptor is proportional to receptor binding affinity. Pseudovirus libraries with Nipah RBP + unmutated F are mixed with ~1% neutralization 'standard' pseudovirus expressing only VSV-G and containing defined barcodes in its genome. Libraries are used to either infect cells with nothing added (the no soluble receptor control) or they are first incubated with soluble receptor at increasing concentrations prior to cell infection. The barcode extraction and sequencing is the same as in A). Counts from the neutralization standard are used to normalize read counts across conditions, followed by quantification of receptor binding by comparing barcode frequencies in the receptor incubation conditions compared to the no receptor control. Mutants that decrease binding to soluble receptor will increase in relative frequency due to decreased neutralization by the receptor. Mutants that increase binding to the receptor will decrease in relative frequency due to being neutralized more. C) Antibody selection overview. A neutralization standard is included for antibody selections, similar to the receptor binding selections. A control condition is also included (the no antibody control), and other conditions use pseudovirus incubated with increasing concentrations of antibody. The barcode extraction and sequencing steps are similar to A) and B). Mutants that increase in relative frequency in the presence of neutralizing antibodies are escape mutations.

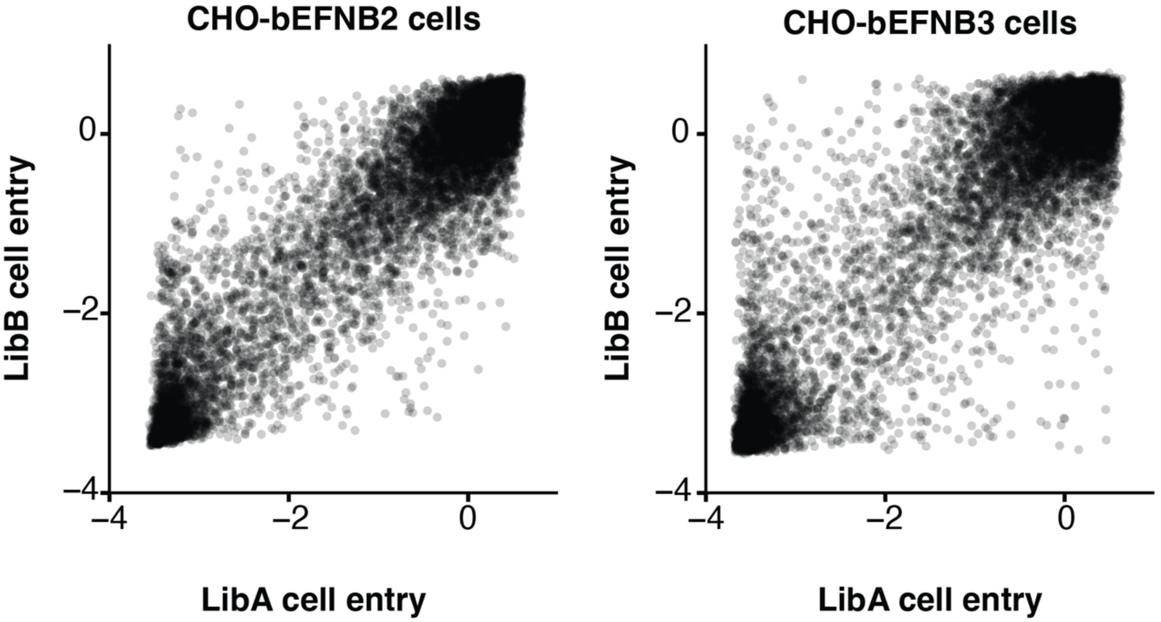


**Supplemental Figure 8. Correlations between effects of RBP mutations on cell entry in CHO-bEFBN2 or CHO-bEFNB3 cells measured using the two independent RBP pseudovirus libraries (LibA and LibB)**. For CHO-bEFNB2 cells, we conducted six independent functional selections using LibA and two using LibB. For CHO-bEFNB3 cells, three functional selections were carried out with LibA and four with LibB. For each library, the value plotted here is the mean measurement for each mutation across the different selections with that library. We only show measurements for mutations present in at least two barcoded RBP variants. The Pearson correlation coefficients (r) were 0.92 for both target cell types.

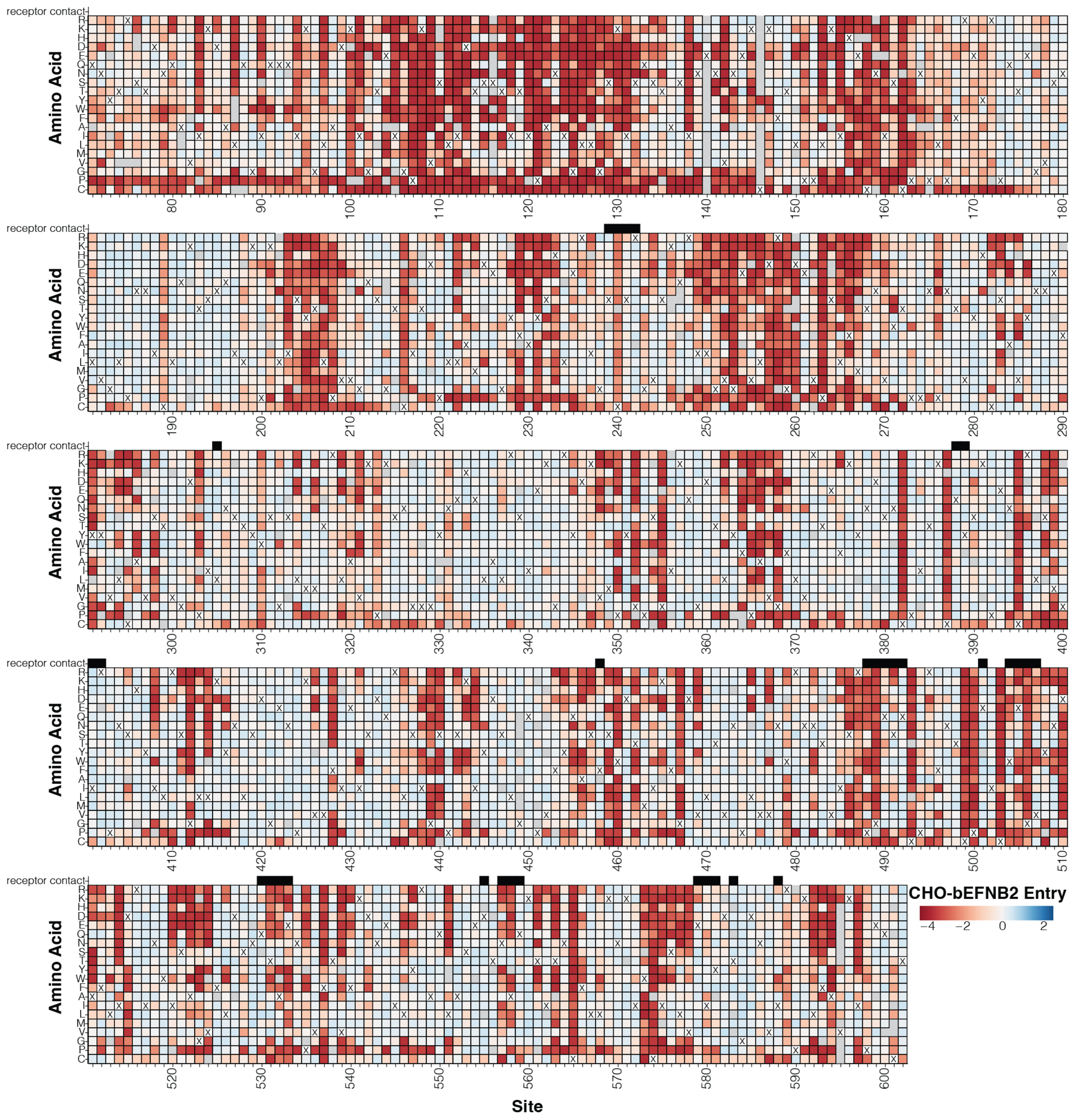


**Supplemental Figure 9. Effects of mutations on entry in CHO-bEFNB2 cells**. This figure is similar to Fig. 1, but shows entry into CHO-bEFNB2 rather than CHO-bEFNB3 cells. Interactive version is available (https://dms-vep.org/Nipah_Malaysia_RBP_DMS/htmls/E2_entry_heatmap.html). Labeling and coloring same as in Fig. 1.

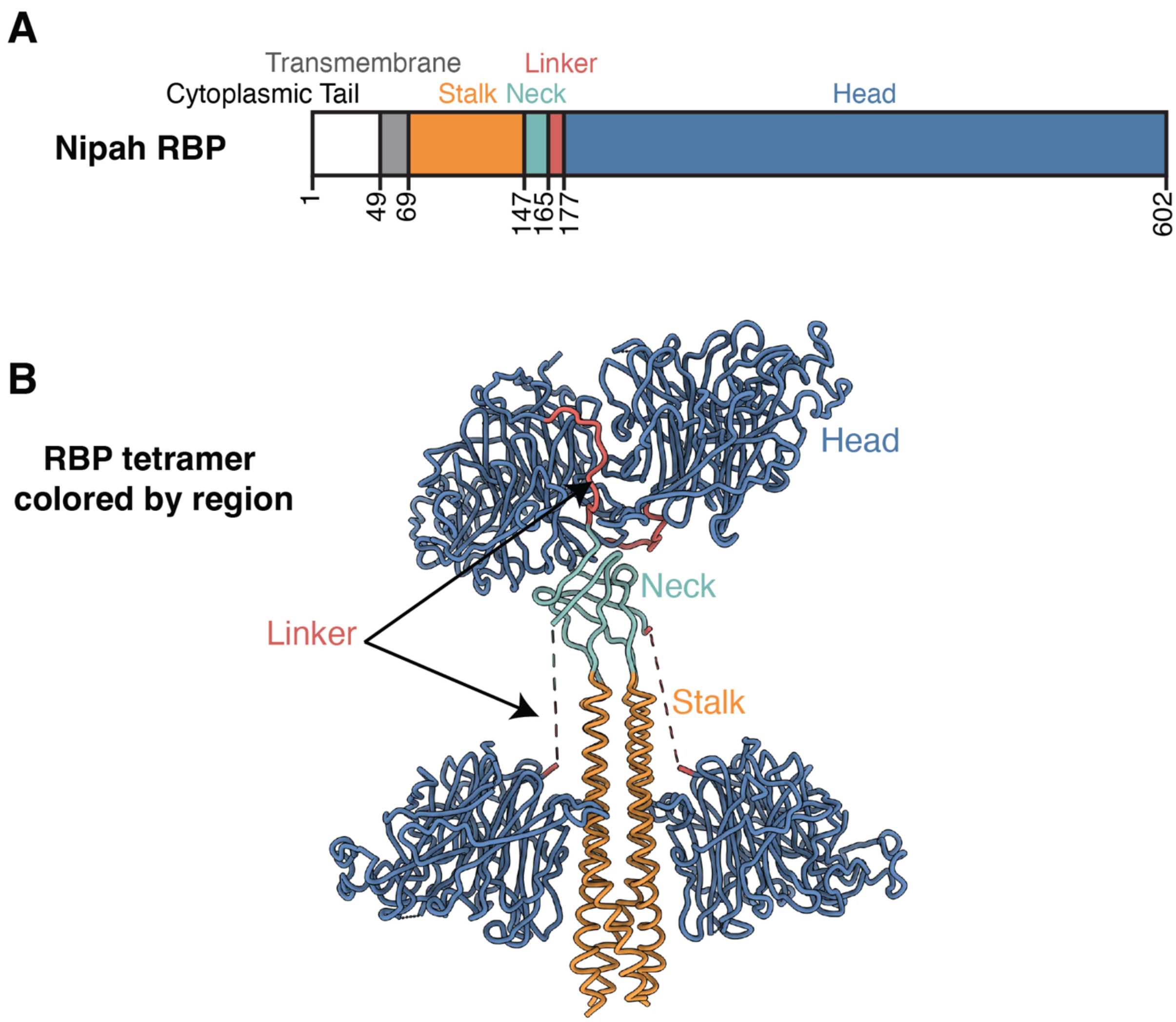


**Supplemental Figure 10. Major regions of Nipah RBP**. A four-helix bundle (stalk) connects to the transmembrane domain that anchors the RBP in the viral membrane. The neck is a β sandwich that has multiple disulfide bonds and is involved in F-triggering. A flexible linker connects the neck with the globular heads. Two of the heads are arranged above the neck (distal), while two are arranged along the stalk towards the viral membrane (proximal). Note the linker region for the proximal heads were not resolved in structure and are indicated with dotted lines. A) Numbering and location of wildtype RBP domains. B) RBP tetramer colored by domain (PDB: 7TXZ,7TY0).

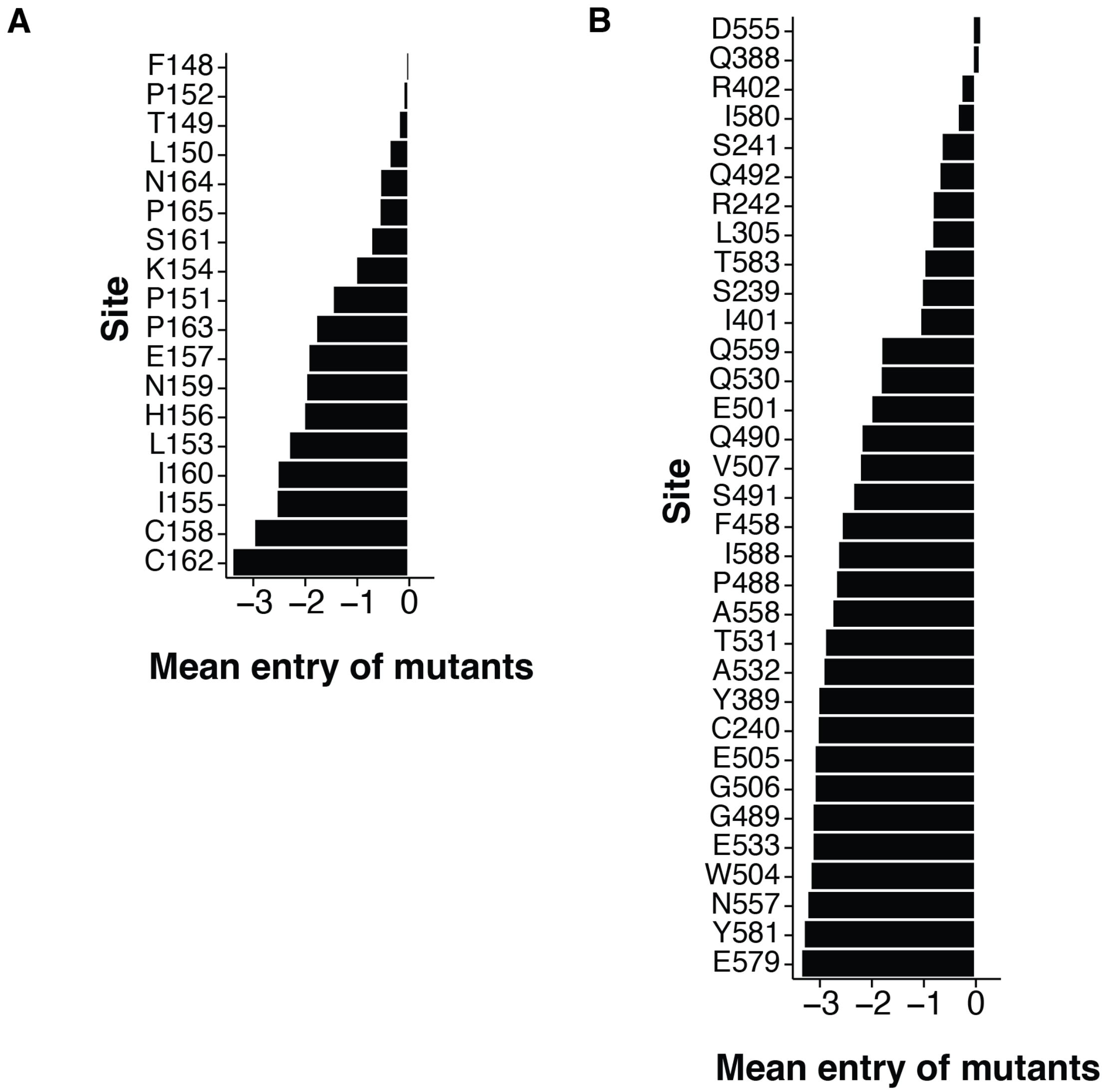


**Supplemental Figure 11. Ranked site-averaged effects of mutations on cell entry in CHO-bEFNB3 cells in the neck and receptor contact residues.** Sites within the neck (A) and receptor contact residues (B) were ranked by the site-averaged effects on cell entry and ordered from least constrained (top) to most (bottom). Most mutations at site 555 are tolerated as EFNB2/3 interacts with the backbone and not the sidechain. E533 is a key contributor to the receptor contact interface, and forms two salt bridges with K60 and K116.

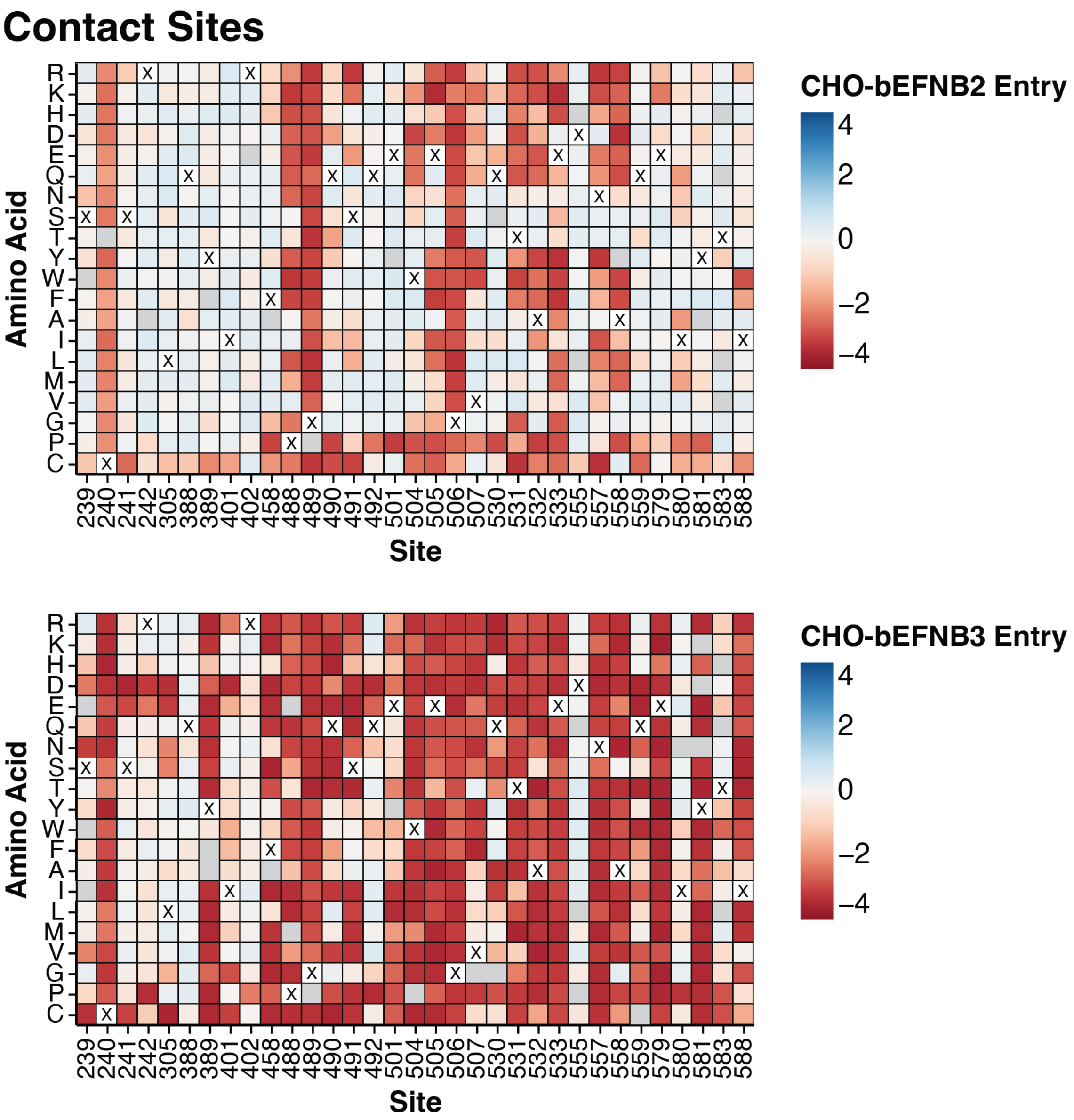


**Supplemental Figure 12. Effects of RBP mutations on entry in CHO-bEFNB2 and CHO-bEFNB3 at receptor contact sites.** Entry scores for mutations in CHO-bEFNB2 cells are shown on top, CHO-bEFNB3 are on bottom. For each mutation, the entry score reflects the cell entry efficiency of a pseudovirus with that RBP mutation relative to the unmutated RBP. Negative values (red) indicate impaired entry, zero (white) indicates no effect, and positive values (blue) indicate improved entry. The unmutated amino-acid in the Malaysia strain RBP at each site is indicated with a 'X'. An interactive version of this figure is available here (https://dms-vep.org/Nipah_Malaysia_RBP_DMS/htmls/combined_entry_binding_contact_heatmaps.html).

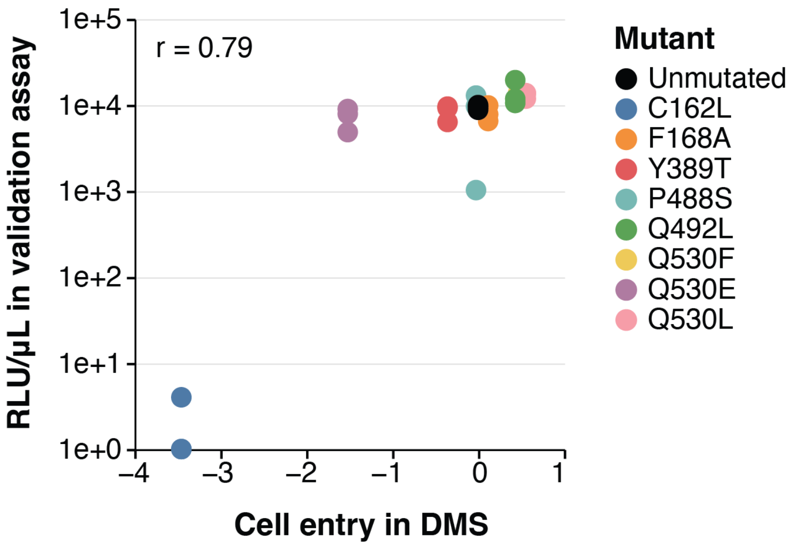


**Supplemental Figure 13. Validations of DMS measurements of mutational effects on cell entry in CHO-bEFNB2 cells with single mutant pseudoviruses compared to unmutated**. The data shown here are similar to that in Fig. 2I except validations were performed with CHO-bEFNB2 rather than CHO-bEFNB3 cells.

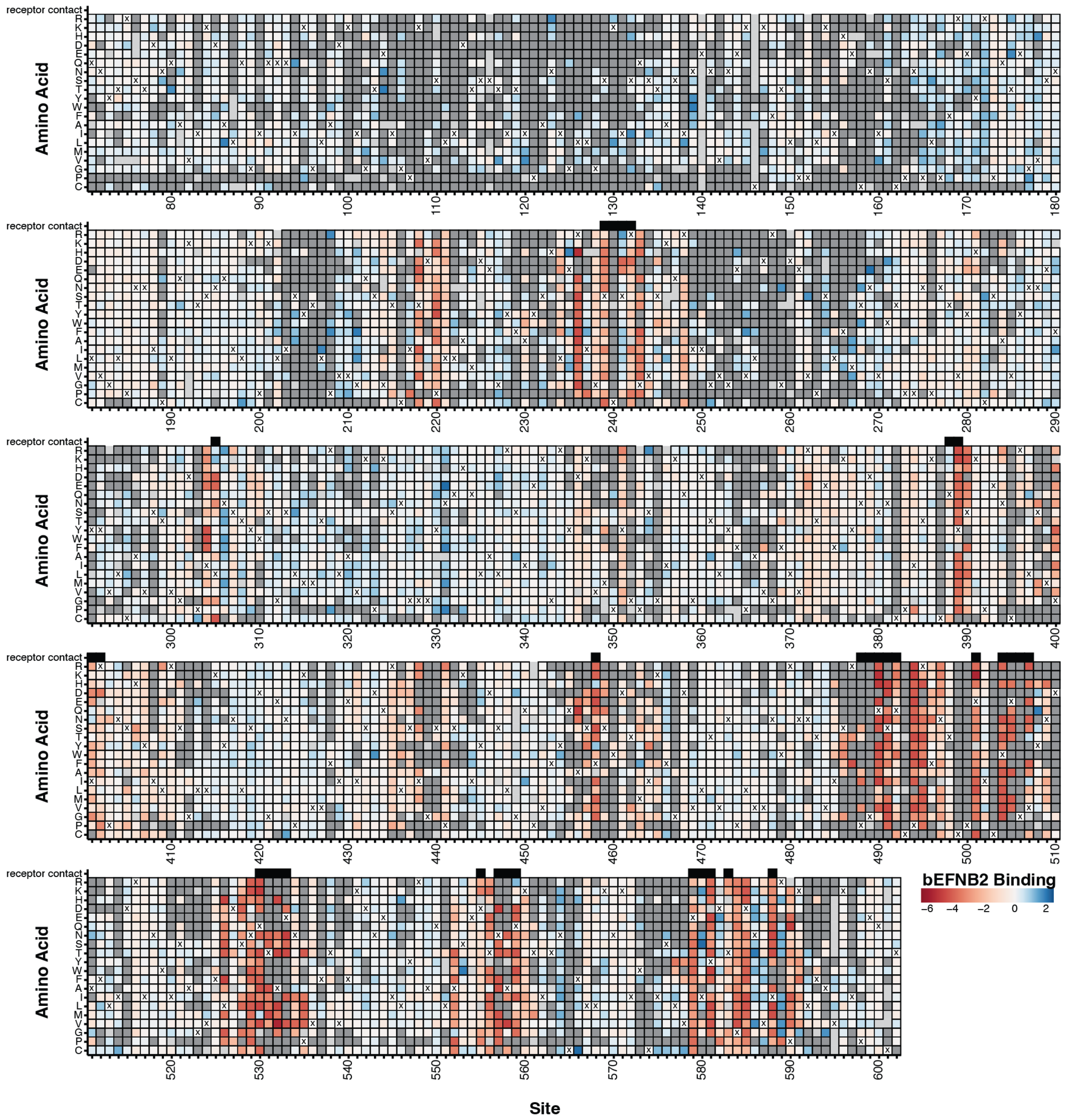


**Supplemental Figure 14. Effects of mutations on binding to monomeric bEFNB2**. Light gray squares are mutants that were missing. Dark gray squares are mutants that were filtered out due to having a low entry score (< -1.5). Receptor contact sites are indicated with a black box above the heat map. The amino-acid identity in the parental Malaysia strain RBP at each site is represented with an 'X'. Mutants that are colored red were neutralized by soluble monomeric bEFNB2 less than the unmutated Nipah RBP (and so have reduced bEFNB2 binding), while mutants colored blue were more neutralized than unmutated (and so have improved bEFNB2 binding). Interactive version can be found here (https://dms-vep.org/Nipah_Malaysia_RBP_DMS/htmls/E2_binding_heatmap.html).

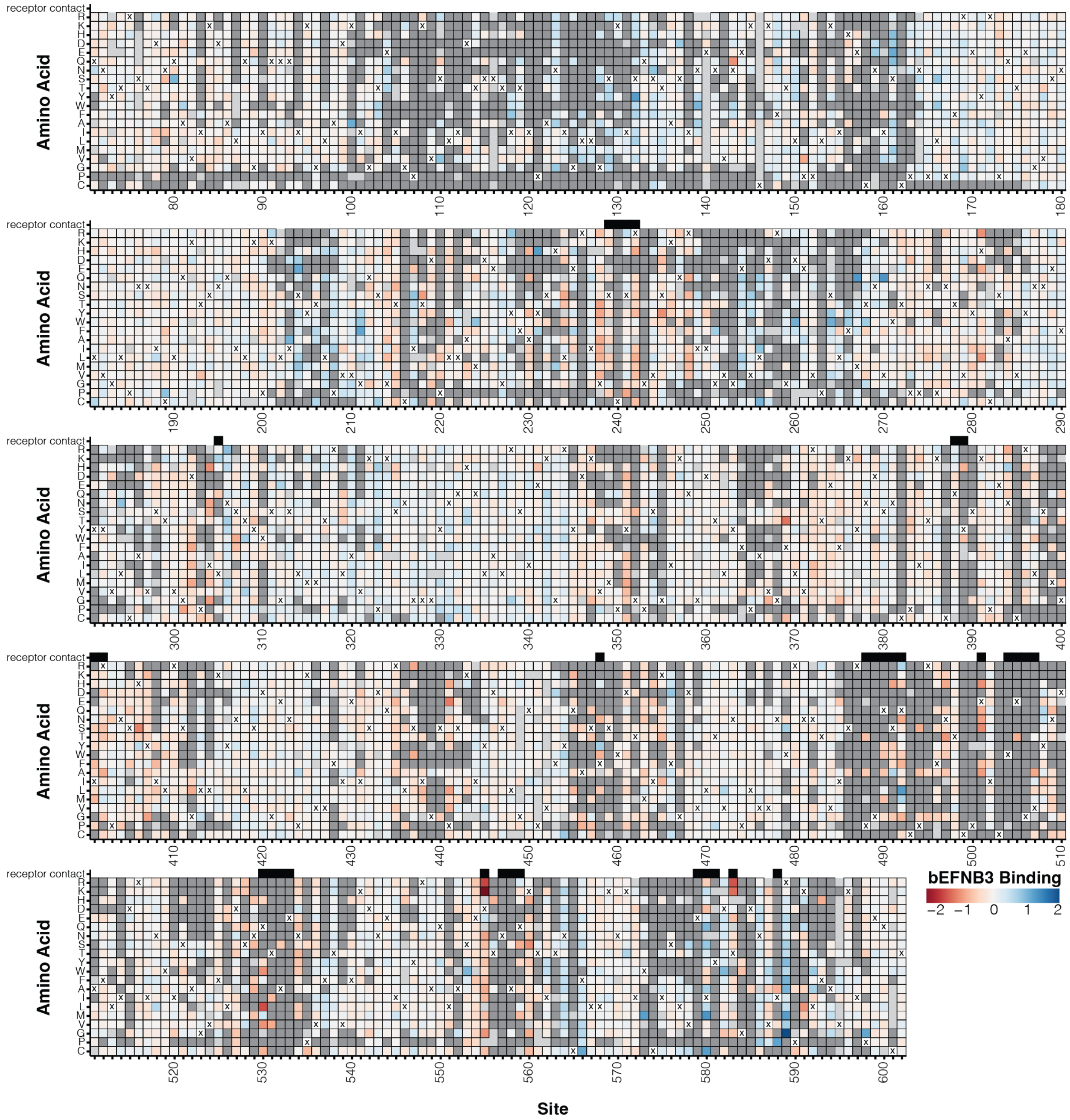


**Supplemental Figure 15. Effects of mutations on binding to dimeric bEFNB3**. See figure legend in Supplemental Figure 14. An interactive heatmap is available here ([https://dms-vep.org/Nipah_Malaysia_RBP_DMS/htmls/E3_binding_heatmap.html](https://dms-vep.org/Nipah_Malaysia_RBP_DMS/htmls/E3_binding_heatmap.html)).

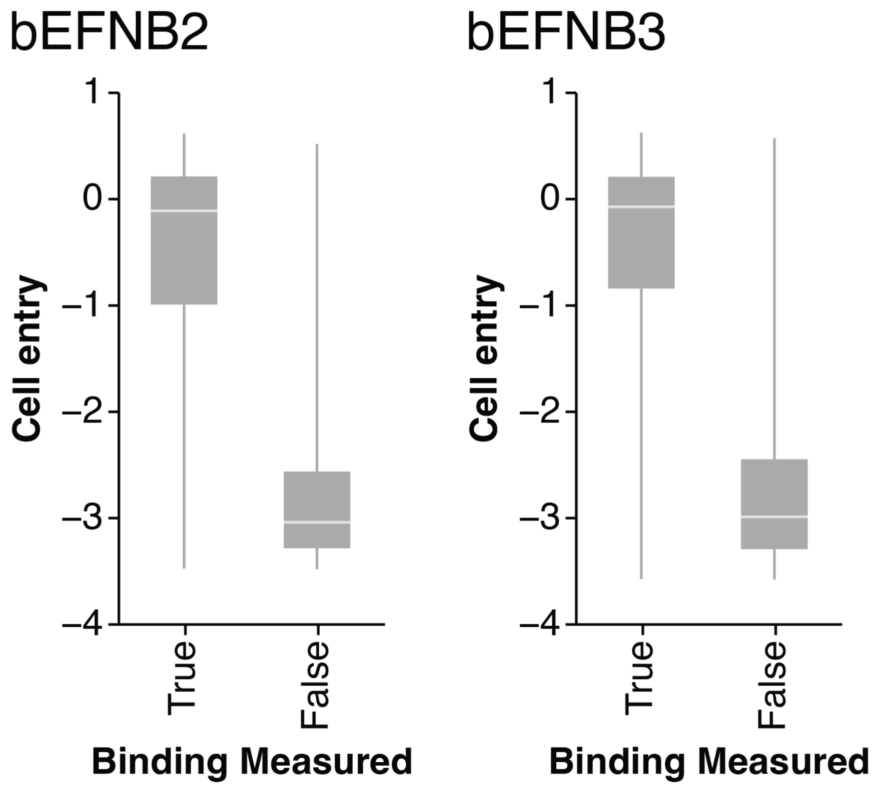


**Supplemental Figure 16. Most mutations for which we could not measure the effect on receptor binding were highly deleterious for cell entry.** Min-max boxplots showing the distribution of cell entry scores for mutations with and without binding measurements. For bEFNB2, we made binding measurements for 8257 of the 9786 mutations with cell entry measurements, and for bEFNB3, we made binding measurements for 7948 of the 9713 mutations with cell entry measurements. The reason that we could not make binding measurements for mutations that are highly deleterious for cell entry is that the experimental approach used to measure binding requires the mutant pseudovirus to still be able to infect cells (fig S7B). However, since the mutations without binding measurements are highly deleterious, they are unlikely to be relevant to actual RBP evolution.

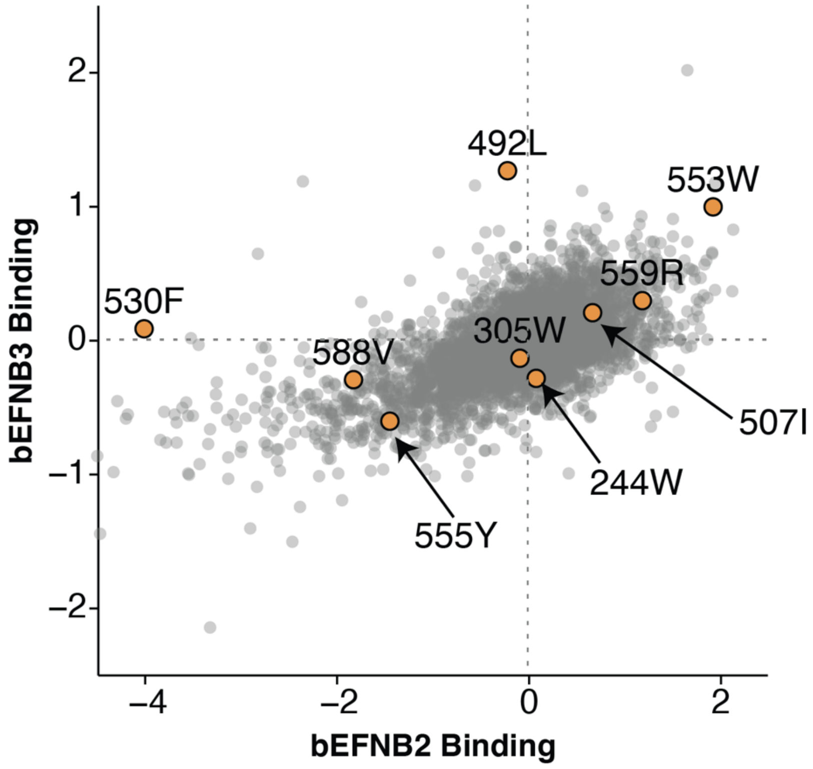


**Supplemental Figure 17. Mutational effects on binding to bEFNB2 and bEFNB3, with mutations picked for validation by biolayer interferometry colored orange.** This plot shows the same data as Fig. 3D, except specific mutations that were validated by biolayer interferometry are colored orange.

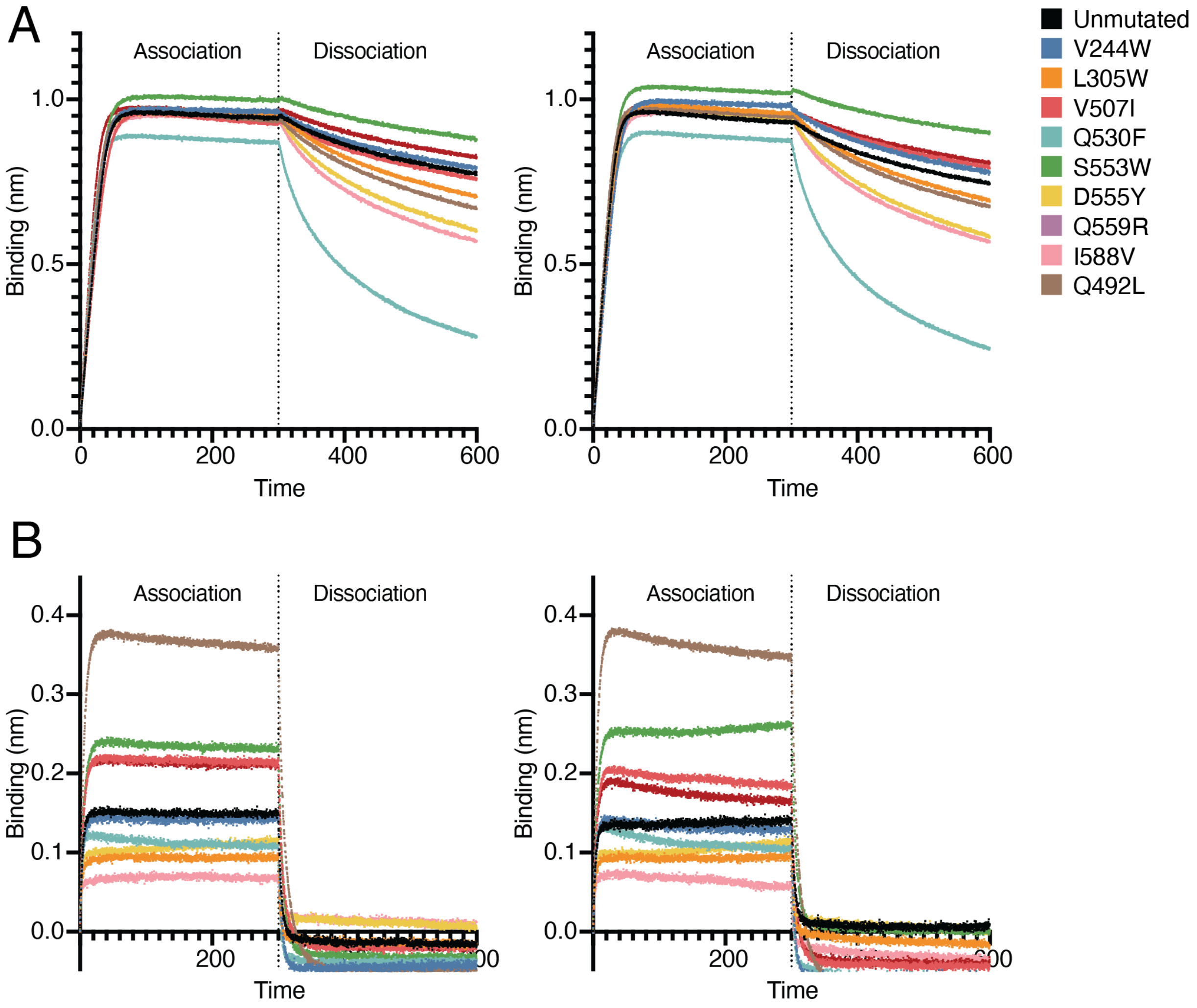


**Supplemental Figure 18. Biolayer interferometry (BLI) sensorgrams for binding of RBP heads to bEFNB2 or dimeric bEFNB3**. BLI sensorgrams from two biological replicates (corresponding to independent batches of proteins) for binding of the unmutated monomeric RBP head or mutants at 50 nM (A) or 100 nM (B) to dimeric bEFNB2 (A) or bEFNB3 (B) immobilized on AHC biosensors. The magnitude of binding was assessed by quantifying the change in total area under the curve (AUC) relative to unmutated RBP for binding of the indicated RBP head domains to immobilized dimeric bEFNB2 or bEFNB3.

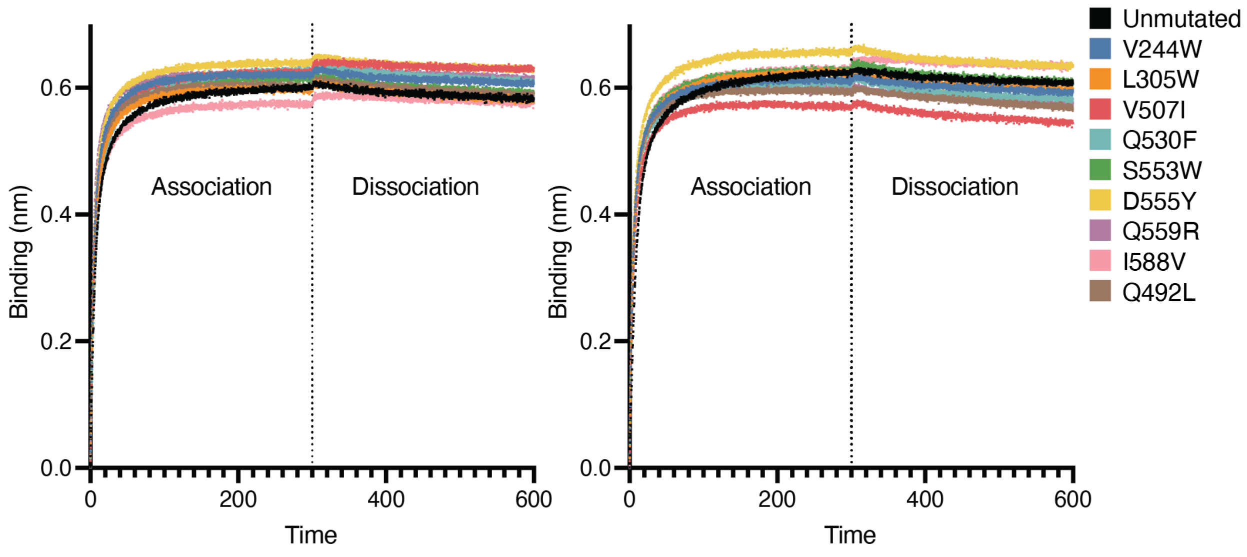


**Supplemental Figure 19. Biolayer interferometry (BLI) sensorgrams for binding of RBP heads to the HENV-32 monoclonal antibody.** BLI binding analysis of purified Nipah virus RBP head wildtype and mutants at 300 nM to HENV-32 IgG immobilized on AHC biosensors. Each sensorgram corresponds to a biological replicate using independently produced batches of proteins.

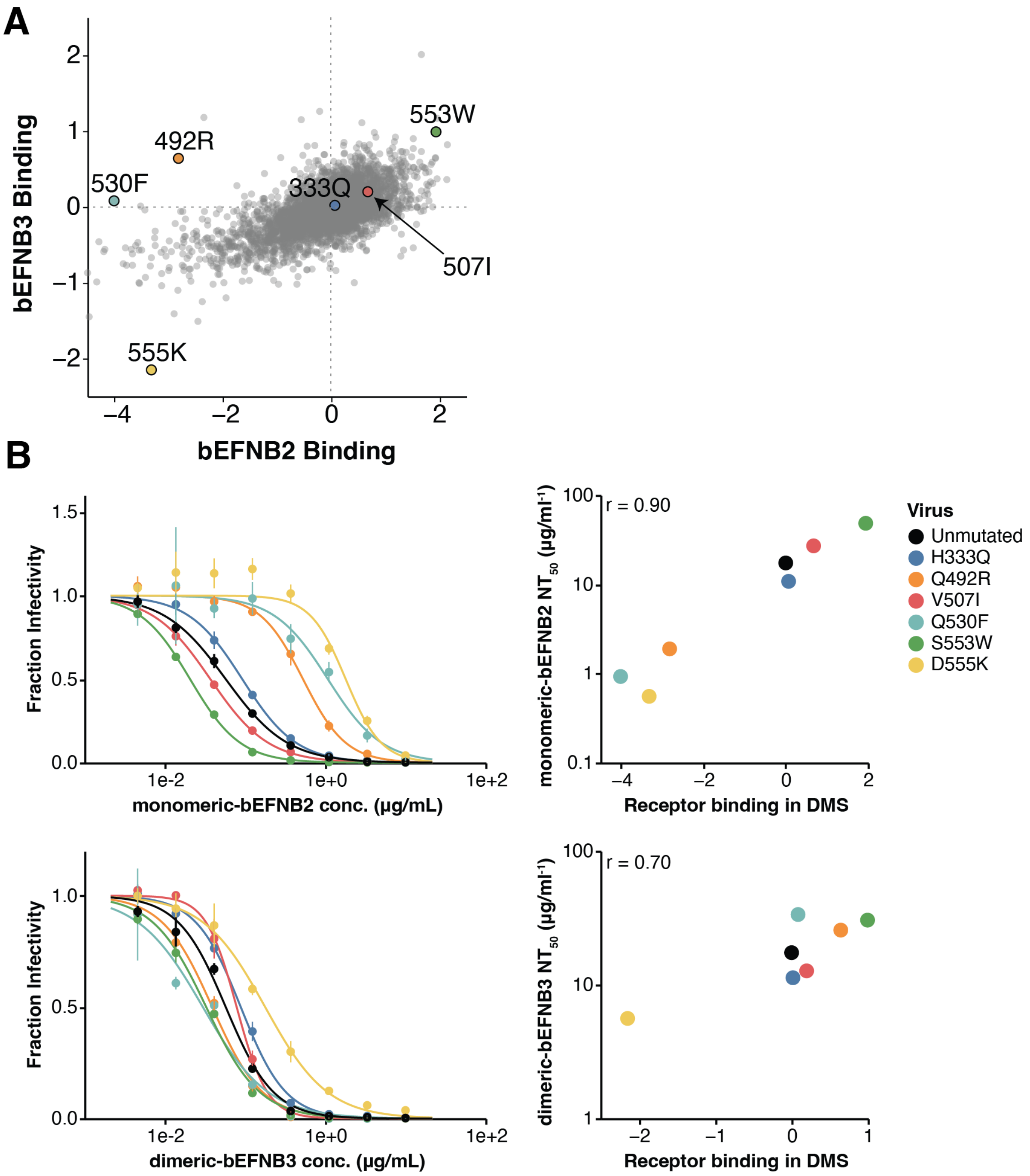


**Supplemental Figure 20. Mutations selected for validations in soluble-receptor neutralization assays and correlations with DMS**. A) Correlation between binding effects for all mutants to bEFNB2 or bEFNB3 (same data as in Fig 3D), but individual mutations that were used for validating mutational effects on neutralization by soluble receptors are colored. B) Neutralization curves of RBP variants with bEFNB2 or bEFNB3 (left). Correlation of $NT_{50}$ with binding values obtained from DMS (right).

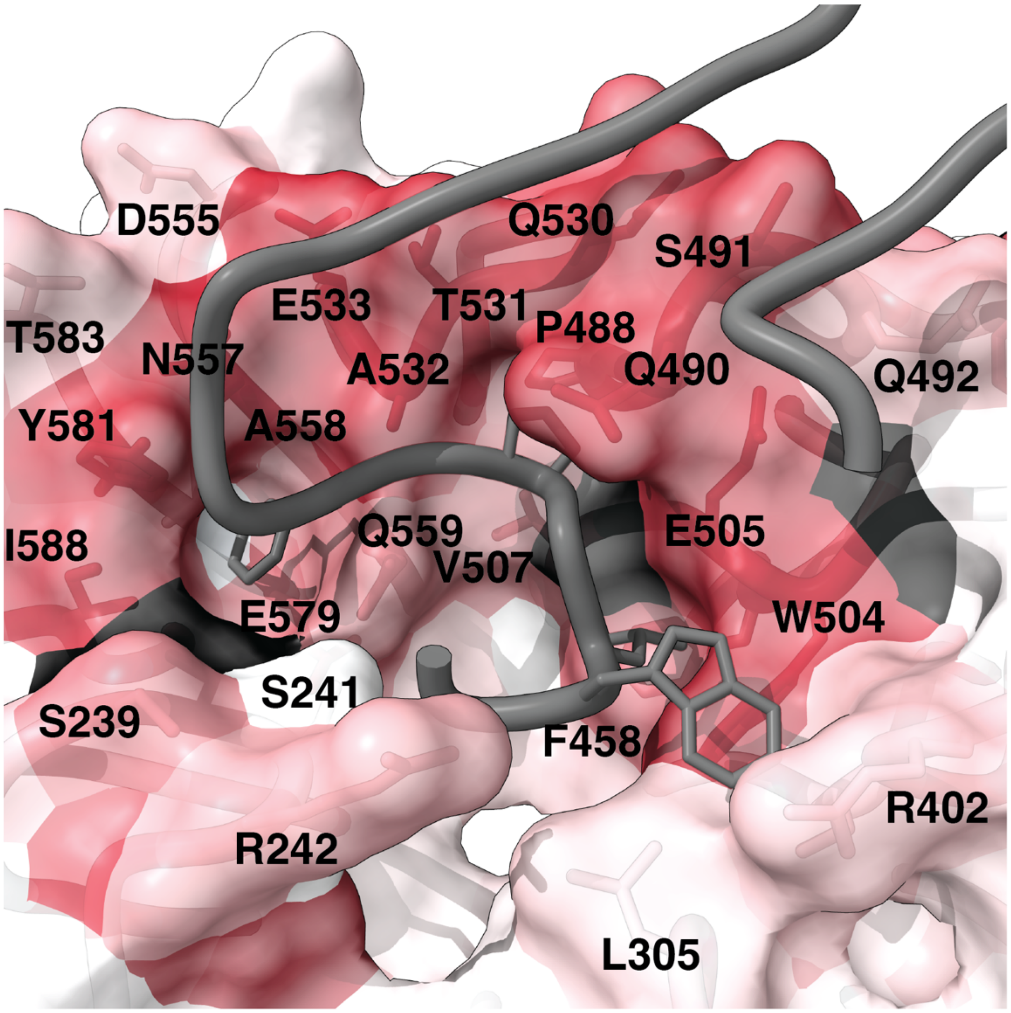


**Supplemental Figure 21. Most mutations in the receptor-binding interface greatly decrease binding affinity.** A) Average effects on bEFNB2 binding of RBP mutations at each site are mapped onto the structure of RBP bound to EFNB2 (PDB: 2VSM). Color scale is the same as in Fig 3C. A portion of the EFNB2 chain is shown (sites 100-123) in dark gray. The four amino-acid residues that insert into hydrophobic pockets on RBPs surface are shown with their side chains (Ephrin sites 120,122,124,125). RBP residues colored black do not have any mutations with binding information, and were filtered out due to low entry scores.

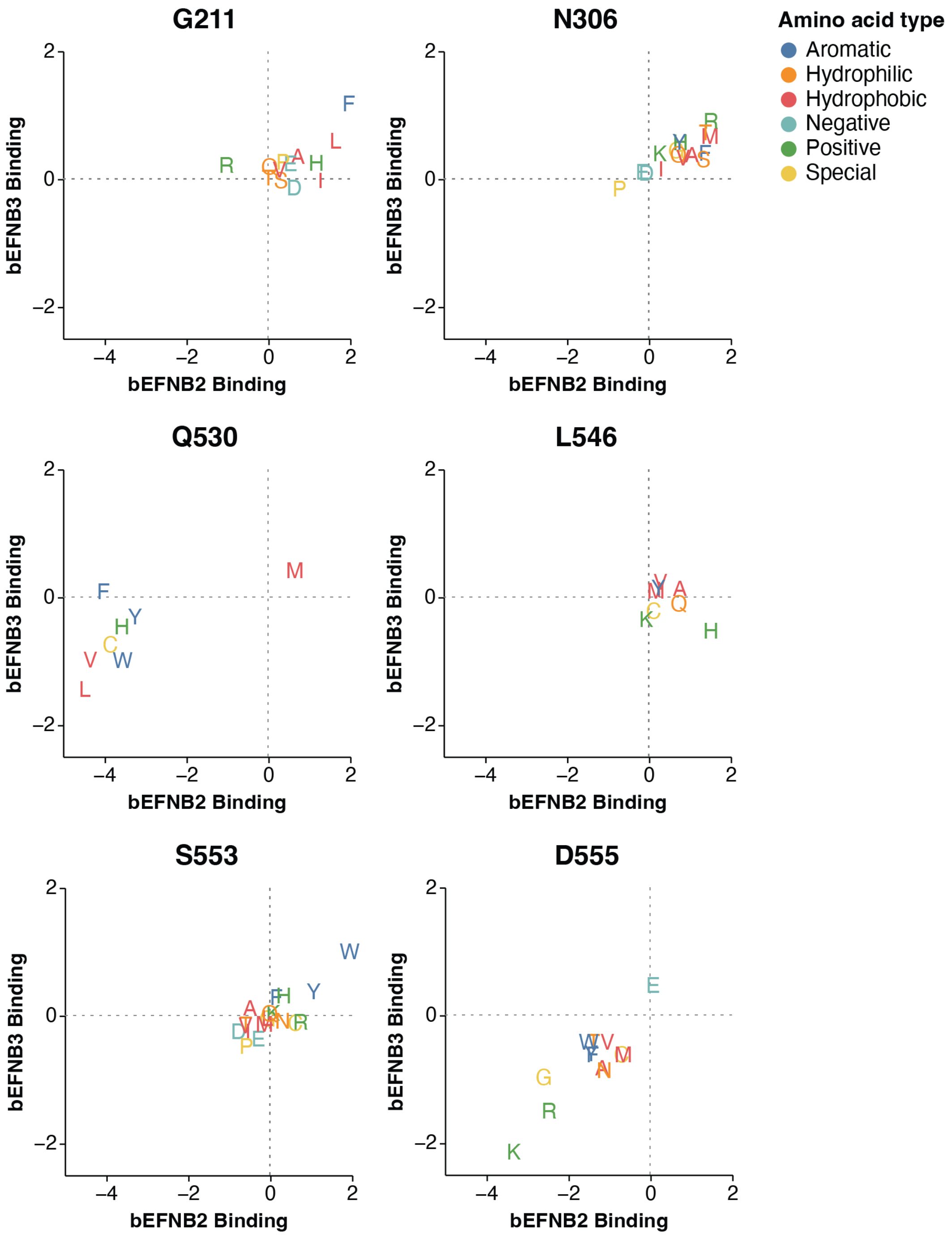


**Supplemental Figure 22. Correlations between effects of mutations on binding to bEFNB2 or bEFNB3 at key sites.** Each panel represents a different site, with the x-axis indicating the binding to bEFNB2 and the y-axis indicating the binding to bEFNB3 as measured in the DMS. The letters indicate the effects of different mutations; the binding score for the unmutated Malaysia RBP is 0 by definition. Sites of interest highlighted in Fig. 3D are shown here. Site 492 is specifically shown in fig S23, and sites of interest in the 580-590 loop are shown in fig S24. Mutations of interest at these sites have large effects on binding to one or both host receptors.

Briefly, G211F increases binding to both bEFNB2 and bEFNB3. Most mutations at site N306 increase binding to bEFNB2/3, likely due to removal of a glycosylation that possibly reduces receptor binding. Q530F greatly decreases bEFNB2 binding, while maintaining bEFNB3 binding (also tested by BLI (Fig. 3B) and pseudovirus neutralization (fig S20). L546H increases bEFNB2 binding while decreasing bEFNB3 binding. S553W/Y increases binding to both bEFNB2/3 (S553W tested by BLI (Fig 3B) and pseudovirus neutralization (fig S20). Finally, substitutions of D555 to positively charged amino acids (K/R), greatly decreases binding to both bEFNB2/3 (D555K tested by pseudovirus neutralization (fig. S18)). An interactive plot for all sites is available at (https://dms-vep.org/Nipah_Malaysia_RBP_DMS/htmls/binding_letter_plot_slider.html).

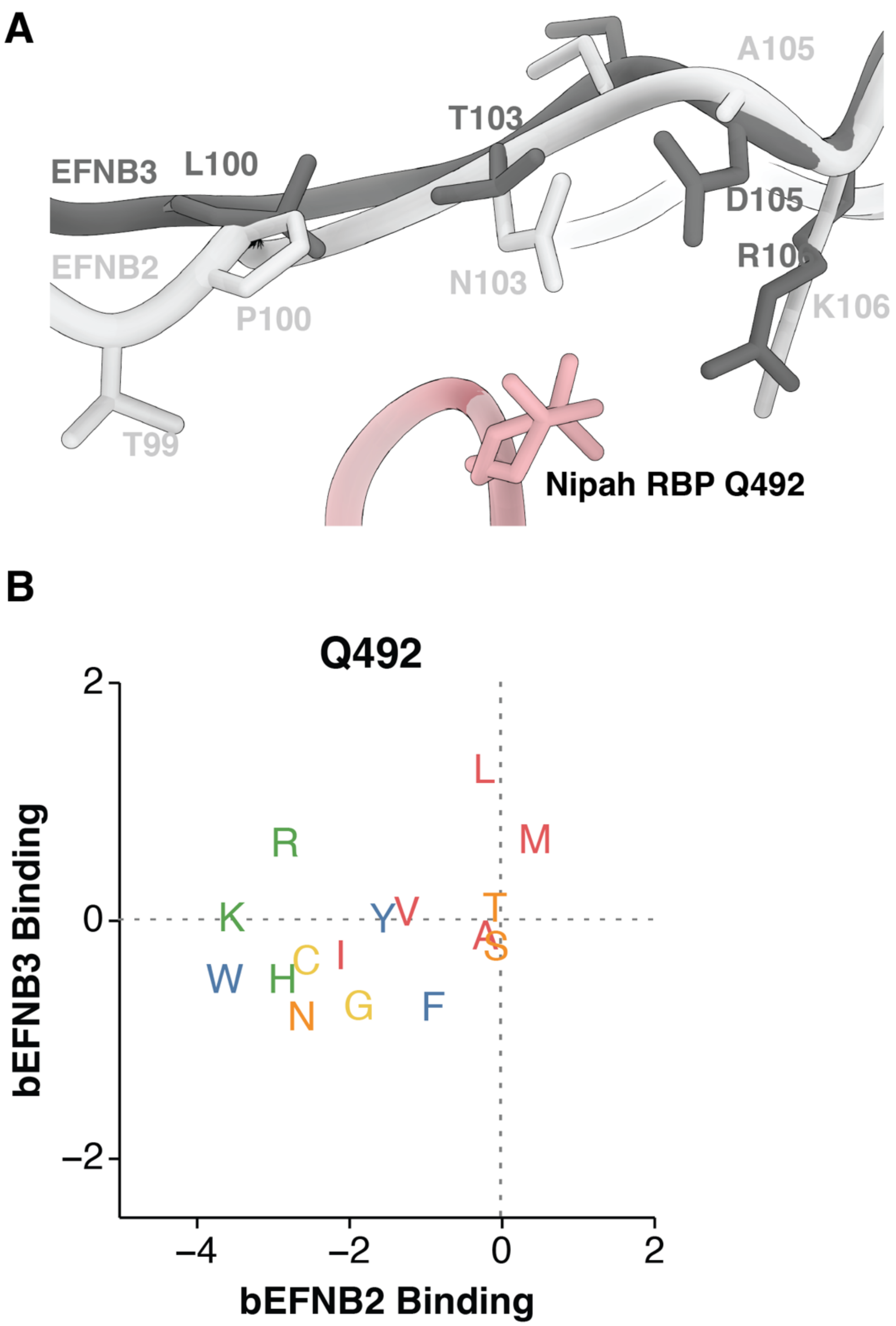


**Supplemental Figure 23. Mutations at RBP site 492 have different effects on binding to bEFNB2 and bEFNB3, and are located near amino acids that are different between each receptor.** A) Superimposed structures of RBP bound to human EFNB2 (light gray; PDB:2VSM) and EFNB3 (dark gray; PDB: 3D12) are shown. Receptor sites that differ between human EFNB2 and EFNB3 are labeled. Site 106 is different between bat and human EFNB2 (human to bat EFNB2 K106R, which changes it to the same residue in EFNB3). B) Correlation between the effects of mutations on binding to bEFNB2 and bEFNB3 at site 492. Mutations to hydrophobic residues such as L and M increase binding to bEFNB3 while maintaining binding to bEFNB2. Mutations to certain positively charged residues (K,R), greatly decrease binding to bEFNB2 while maintaining or even increasing binding to bEFNB3. The increased binding of Q492R/K mutations is likely from a salt bridge formed with D105 in bEFNB3. Q492L affinity was directly measured by BLI (Fig. 3B). Q492R was tested by pseudovirus neutralization (fig S20).

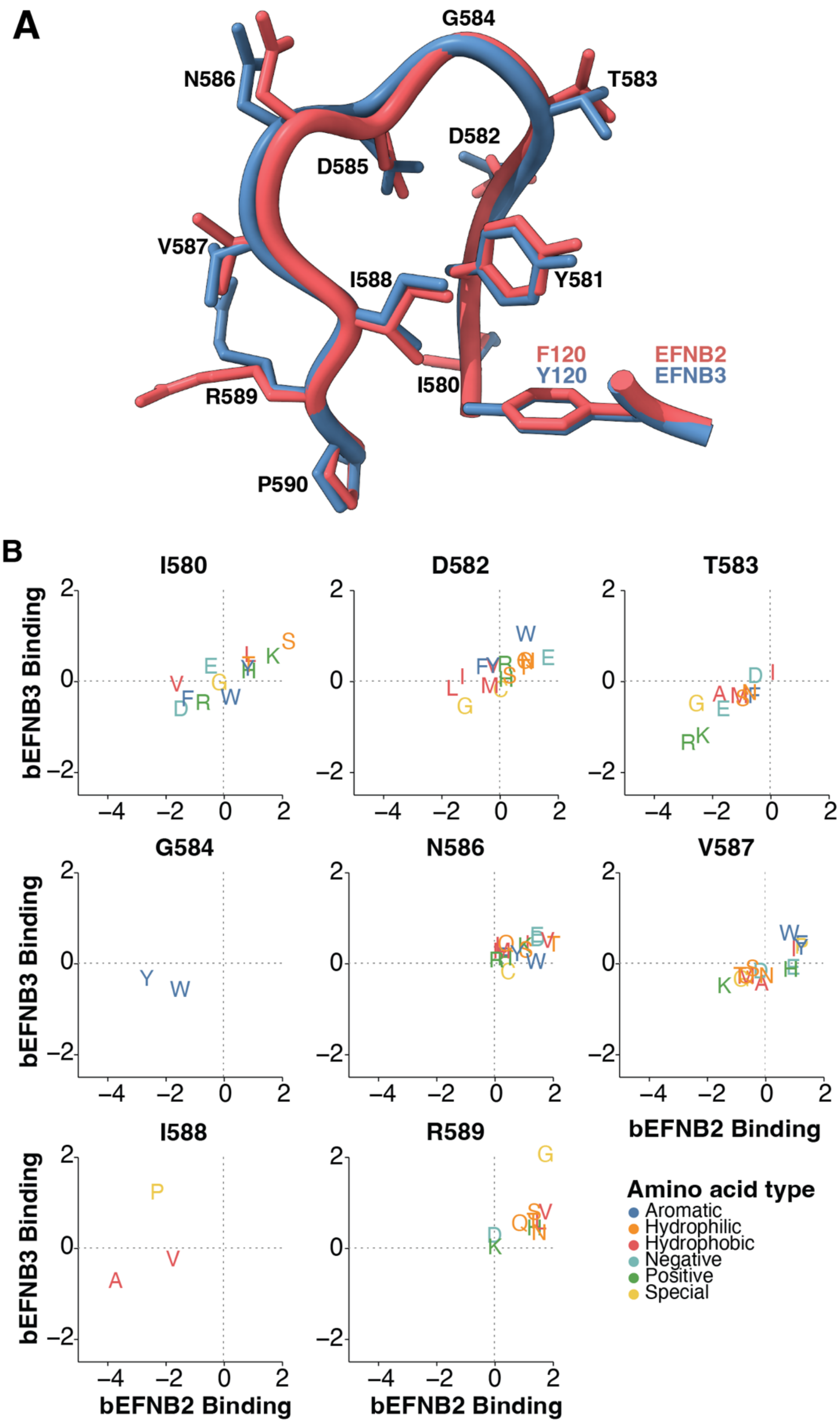


**Supplemental Figure 24. Effects of mutations on binding to either bEFNB2 or bEFNB3 at RBP sites 580-590**. Certain sites in RBP loop 580-590 have mutations with large effects on binding to bEFNB2 or bEFNB3. A) Structures of RBPs bound to either EFNB2 (colored red;

PDB: 2VSM) or EFNB3 (colored blue; PDB:3D12). Ephrin site 120, which is a F in EFNB2 and a Y in EFNB3 is shown on the right. B) Differences in effects of individual mutations on binding to either bEFNB2 or bEFNB3 at sites within the 580-590 loop. Only sites with mutations that affect binding are shown. Sites with mutations that increase binding to both bEFNB2 and bEFNB3 have side chains pointed away from the receptor contact residue (sites 580, 582, 586, 587, 589).

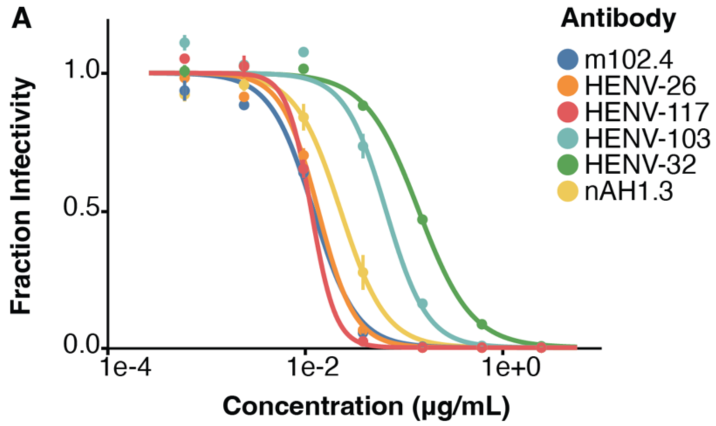


B

| Antibody | IC50 (ng/mL) | Epitope |
| --- | --- | --- |
| m102.4 | 12.3 | Receptor binding face |
| HENV-26 | 13.7 | Receptor binding face |
| HENV-117 | 11.7 | Receptor binding face |
| HENV-103 | 66.7 | Dimerization face |
| HENV-32 | 143.0 | Dimerization face |
| nAH1.3 | 23.1 | Distal face |

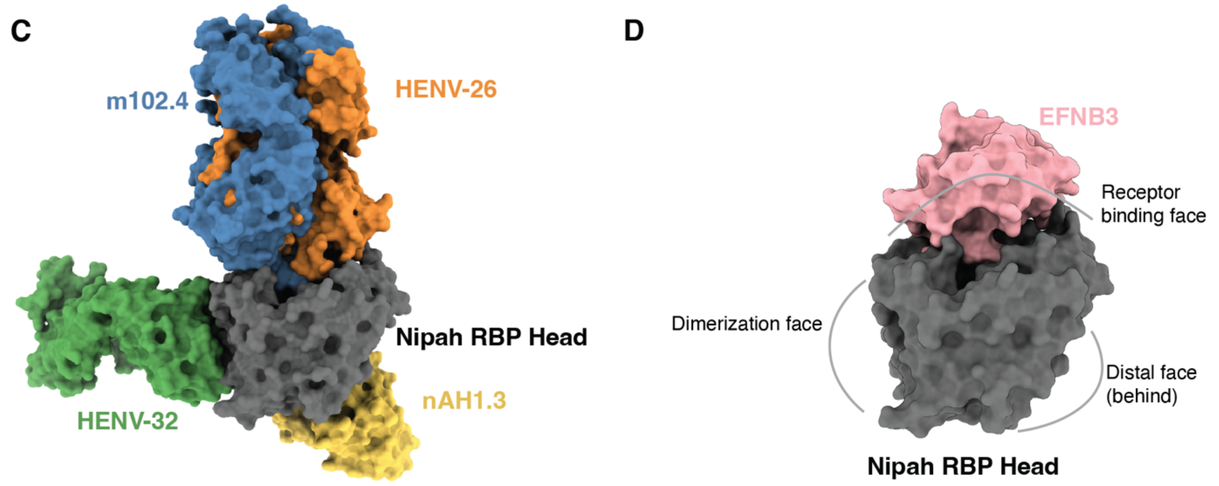


**Supplemental Figure 25. Neutralization of Nipah pseudovirus by antibodies.** A) Neutralization curves of unmutated pseudovirus with each antibody. Fraction infectivity was determined by incubating pseudovirus containing a luciferase reporter with eight different concentrations of antibody. 48 hours after infection, luciferase signal was determined on a plate reader and fraction infectivity was calculated by normalizing to conditions without antibody. All measurements were done in duplicate. B) Table of the $IC_{50}$ in ng/mL for each antibody, as estimated from the neutralization curves in A. C) Antibody binding locations on RBP's head for the four antibodies with structural data. D) Location of epitopes on RBP's head with respect to EFNB3. RBP's head is shown in the same orientation as C. The PDB ID for m102.4 was 6CMG, HENV-26 6VY5, HENV-32 6VY4, EFNB3 3D12, and nAH1.3 was 7TXZ. Note: for 6CMG, the structure was solved with Hendra RBP and m102.3, which is nearly identical to m102.4. For HENV-32, the structure was solved with Hendra RBP.

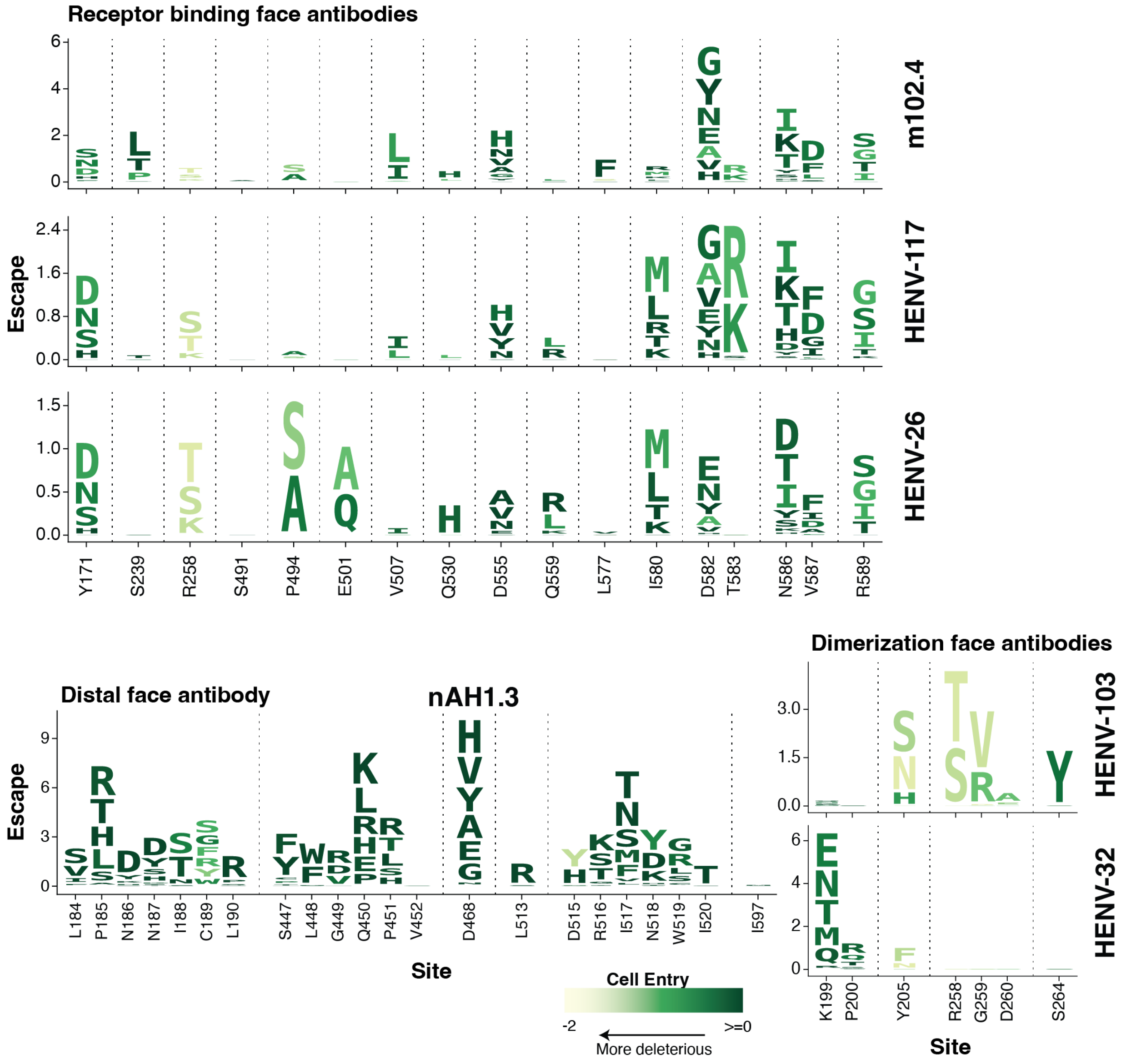


**Supplemental Figure 26. Logo plots similar to those in Figure 4 but showing only amino-acid mutations that are accessible from a single-nucleotide change in the Nipah Malaysia reference strain (Genbank accession NC_002728.1)**. Refer to the legend in Fig 4A for information about the scale and coloring scheme used.

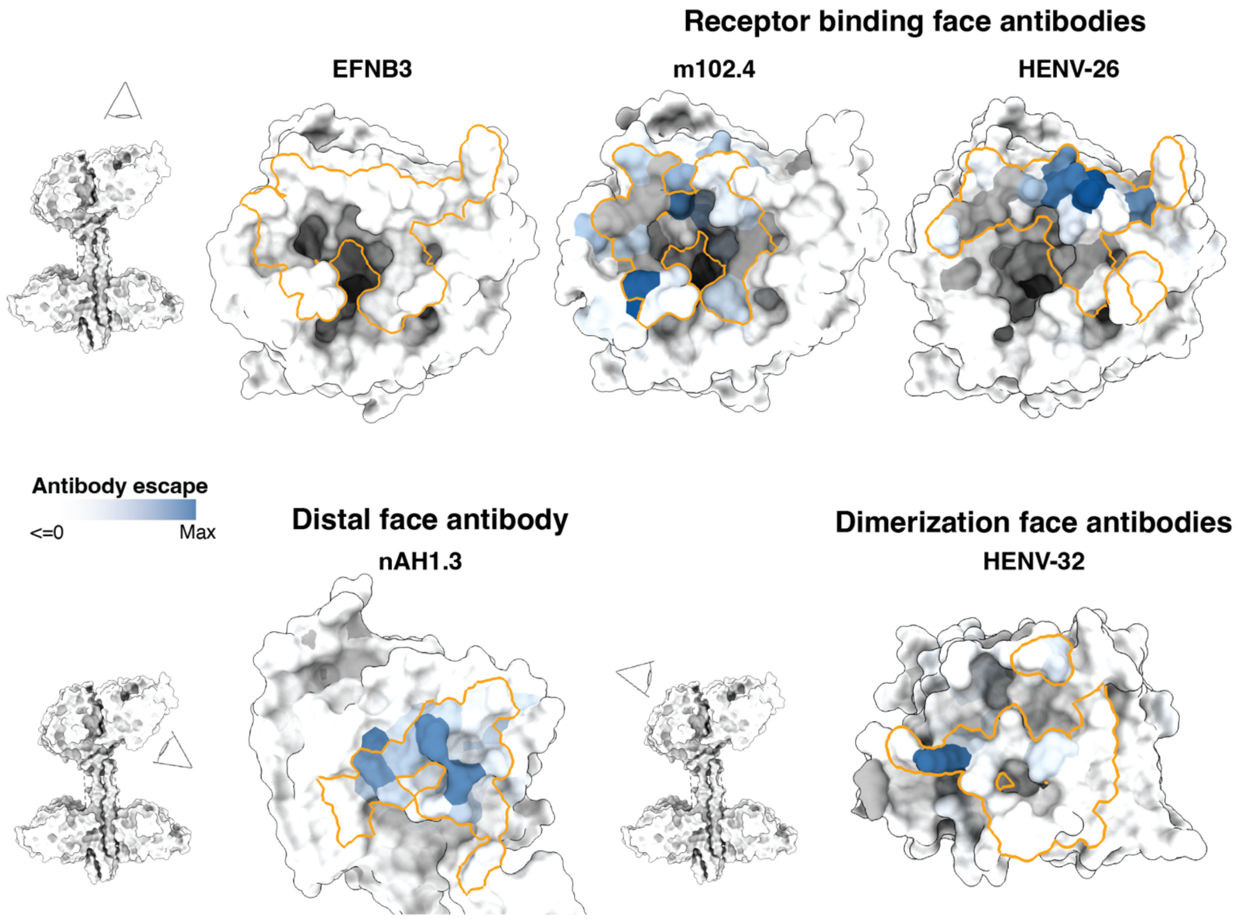


**Supplemental Figure 27. Sites of escape are located in the antibody binding footprints**. Structures of RBP's head bound by each antibody for the four antibodies with structures available. Orange lines outline where an antibody binds, calculated as all RBP sites within 4 angstroms of an antibody residue. Site-average escape as measured in DMS are colored with a white-to-blue color scale (blue = higher escape), with key RBP site numbers indicated in text. Sites with no mutant data (filtered out due to having low entry) are colored light gray. The relative view of each RBP is shown with an associated eye logo and RBP tetramer. The EFNB3 binding area is shown for reference. The PDB ID for m102.4 was 6CMG, HENV-26 6VY5, HENV-32 6VY4, EFNB3 3D12, and nAH1.3 was 7TXZ. Note: for 6CMG, the structure was solved with Hendra RBP and m102.3, which is nearly identical to m102.4. For 6VY4, Hendra RBP was solved with HENV-32.

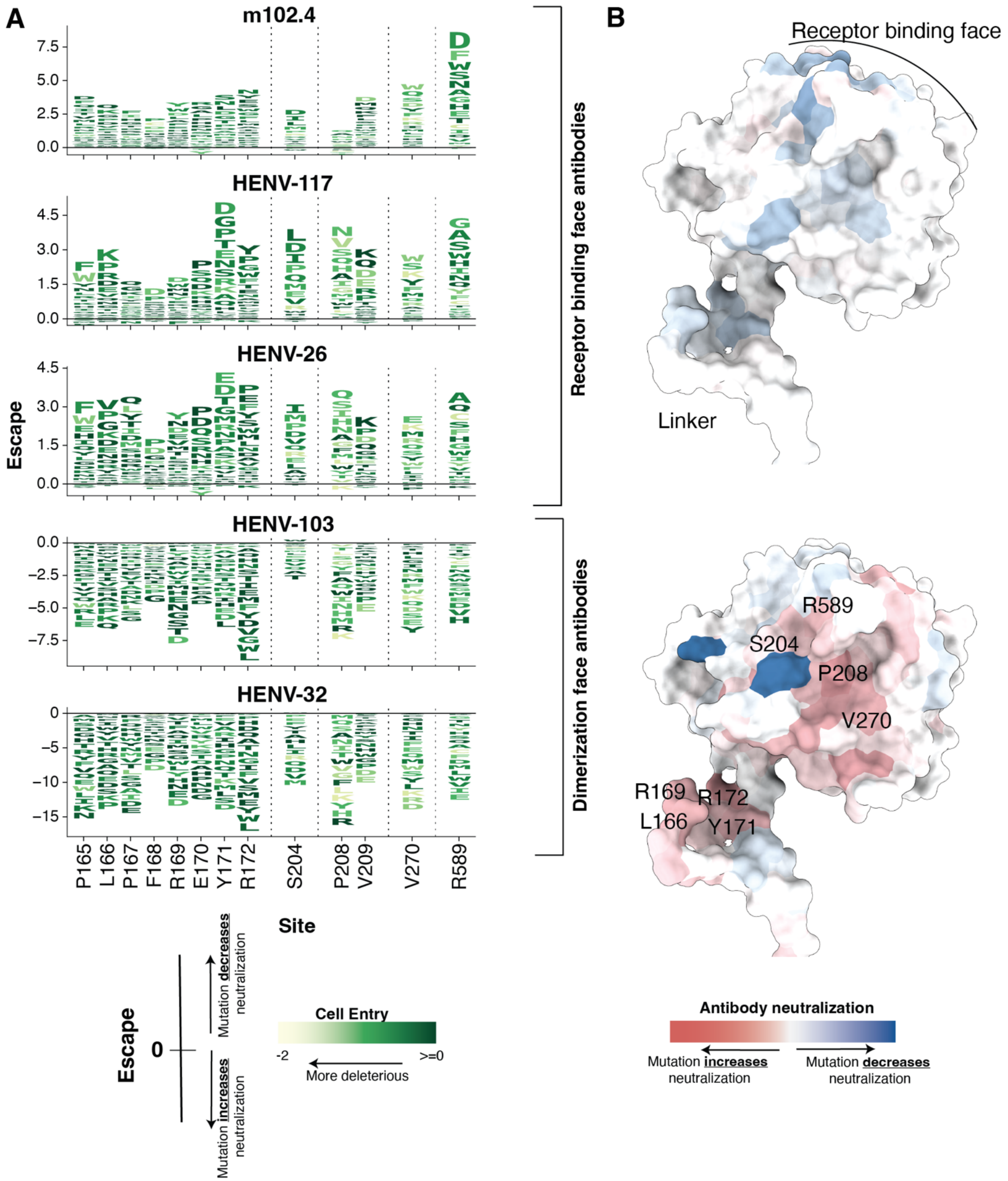


**Supplemental Figure 28. Opposing effects of mutations on neutralization between receptor binding and dimerization face targeting antibodies.** A) Effect of mutations on antibody neutralization at specific sites. The same scale and coloring scheme is used in Fig. 4A, except mutations that also increase neutralization are shown (negative letters below the center line). The height of each letter is proportional to the change in neutralization compared to the unmutated sequence. Mutations with negative scores increase neutralization by that antibody, while mutations with positive scores decrease neutralization. Note that there are escape mutations for HENV-103 and HENV-32 (Fig. 4A), they just do not fall in the 13 sites shown here. B) Site-averaged effects of mutations on antibody neutralization at RBP escape sites. Averages between the three receptor binding face antibodies (top) and the two dimerization face antibodies (bottom) are shown. Sites with mutations that tend to increase antibody neutralization are colored red, those that decrease neutralization are colored blue. Selected sites from (A) are shown on the bottom structure.

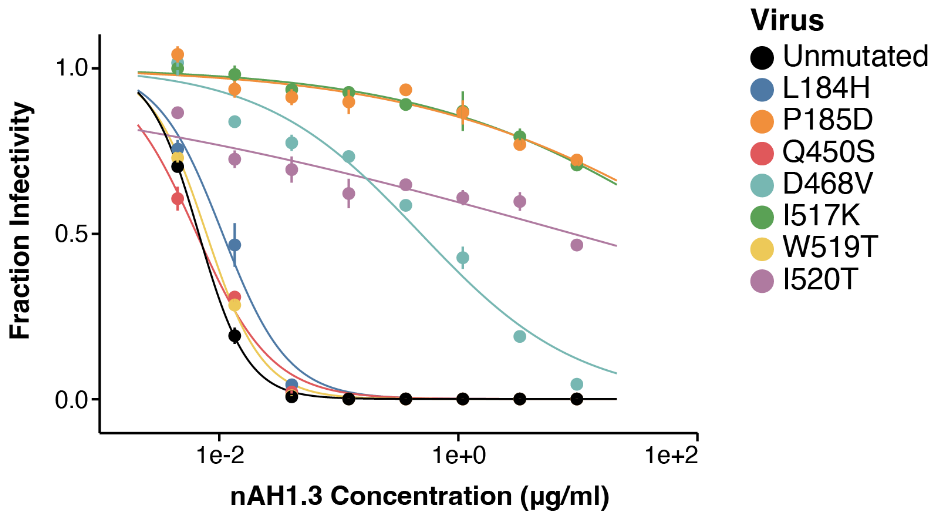


**Supplemental Figure 29. Neutralization curves of different RBP pseudoviruses used to validate measurements of antibody escape from DMS.** Neutralization curves of unmutated and single mutant pseudoviruses against the antibody nAH1.3. Eight different concentrations of antibody were used, and fraction infectivity was determined by comparing the luciferase signal at each concentration to the signal in the absence of antibody. Assays were performed in duplicate. The $IC_{50}$ values from these neutralizations were used to make the correlation in Fig 4D.

| Antibody | Escape | Virus | Type | Reference |
|---|---|---|---|---|
| m102.4 | V507I | Nipah | Authentic virus | Borisevich 2016 |
| m102.4 | D582N | Hendra | Authentic virus | Borisevich 2016 |
| m102.4 | V507I | Nipah | Authentic virus | Xu 2013 |
| m102.4 | D582N | Hendra | Authentic virus | Xu 2013 |
| m102.4 | D582N | Hendra | Cedar chimera | Wang 2022 |
| nAH1.3 | Q450K | Nipah | Authentic virus | Borisevich 2016 |
| nAH1.3 | R516K | Nipah | Authentic virus | Borisevich 2016 |
| nAH1.3 | I520T | Nipah | Cedar chimera | Wang 2022 |
| nAH1.3 | T117A, N186D | Hendra | Cedar chimera | Wang 2022 |
| nAH1.3/m102.4 cocktail | T117A, N186D, T507I | Hendra | Cedar chimera | Wang 2022 |

**Supplemental Figure 30. Antibody escape mutations previously identified in low throughput assays.** Previously described RBP escape mutations from the literature (*31*, *32*, *45*). Includes information about which antibody was used, and the reference for the study. A Cedar chimera is a GFP-encoding replicating Cedar virus (a non-pathogenic henipavirus) with the glycoproteins swapped to Nipah RBP/F.
